# Cellulose synthase-like D (CSLD) proteins move in the plasma membrane and their targeting to cell tips, but not cell plates, depends on the actin cytoskeleton

**DOI:** 10.1101/2021.08.19.457018

**Authors:** Shu-Zon Wu, Arielle M. Chaves, Rongrong Li, Magdalena Bezanilla, Alison W. Roberts

## Abstract

Cellulose Synthase-Like D (CSLD) proteins are implicated in cell wall remodeling during tip growth and cell division in plants, and are known to generate β-1,4-glucan. It is unknown whether they form complexes and move in the plasma membrane like members of the Cellulose Synthase (CESA) family. We used the genetically tractable moss *Physcomitrium patens*, which has a filamentous protonemal stage that undergoes both tip growth and cell division and is amenable to high resolution live cell imaging, to investigate CSLD function and intracellular trafficking. *CSLD2* and *CSLD6* are highly expressed in gametophores and are redundantly required for gametophore cellular patterning. Live cell imaging revealed that CSLD6 is also expressed in protonemata where it moves in the plasma membrane and localizes to cell plates and cell tips. Notably, delivery to the apical plasma membrane, but not the cell plate, depends on actin. By comparing the behavior of endogenously tagged CSLD6 and CESA10, we discovered that CSLD6 movements in the plasma membrane were significantly faster, shorter in duration and less linear than CESA10 movements and were insensitive to the cellulose synthesis inhibitor isoxaben. These data suggest that CSLD6 and CESA10 function within different structures and may thus produce structurally distinct cellulose microfibrils.

## Introduction

Cellulose, a major component of plant cell walls, plays essential biological roles for plants and serves important practical roles for people. For example, cellulose helps regulate plant morphogenesis (Bidhendi and Geitmann 2016) and is being considered as a renewable and carbon-neutral replacement for fossil fuels (Carroll and Somerville 2009). Unique among land plant cell wall components, cellulose is synthesized as microfibrils, which consist of β-1,4-glucan chains joined laterally through hydrogen bonding. The mechanical properties of cellulose microfibrils depend on their cross-sectional dimensions and number of glucan chains, which in turn depend on the structure of Cellulose Synthase Complexes (CSCs) responsible for both glucan polymerization and microfibril assembly (Tsekos 1999; Brown 1996). The rosette-type CSCs of land plants are composed of six particles, each containing three Cellulose Synthase (CESA) catalytic subunits (Nixon et al. 2016; Vandavasi et al. 2016; Kimura et al. 1999; Pear et al. 1996; Mueller and Brown 1980), and produce microfibrils consisting of 18 glucan chains (Jarvis 2018; Oehme et al. 2015; Kubicki et al. 2018).

The CSC was first conceptualized in the ‘ordered granule hypothesis’ (Preston 1964) and posited to move in the plasma membrane (Brown and Montezinos 1976) propelled by the force of cellulose crystallization (Herth 1980; Roberts et al. 1982; Diotallevi and Mulder 2007). Live cell imaging of a YFP-CESA6 translational fusion was a critical advance that provided direct evidence of CSCs movement (Paredez et al. 2006) and enabled visualization and quantification of CSC behavior (McFarlane et al. 2014), identification of cellular compartments involved in CESA trafficking (Crowell et al. 2009; Gutierrez et al. 2009), and characterization of the role of actin and myosin in CSC delivery to the plasma membrane (Sampathkumar et al. 2013; Zhang et al. 2019). Cellulose biosynthesis inhibitors (CBIs) are useful tools for acute inhibition of CESAs (Tateno et al. 2016). Live cell imaging in Arabidopsis has revealed modes of action, which differ among CBI classes (Brabham and Debolt 2012). For example isoxaben clears CESAs from the plasma membrane (Paredez et al. 2006), whereas 2,6-dichlorobenzonitrile (DCB) promotes accumulation of CESAs in the plasma membrane while blocking their movement (DeBolt et al. 2007).

While cellulose biosynthesis by CESAs has been well studied, recent evidence suggests that related Cellulose Synthase-like D proteins (CSLDs) also synthesize cellulose-like β-1,4-glucan (Park et al. 2011; Yang et al. 2020; Hu et al. 2019). Like CESAs, functional CSLDs reside in the plasma membrane (Park et al. 2011) and they synthesize β-1,4-glucan *in vitro* (Yang et al. 2020). However, it is not known whether the *in vivo* product of CSLDs is microfbrillar. In Arabidopsis and other plants, specific CSLDs are required for the development of pollen tubes (Doblin et al. 2001; Bernal et al. 2008) and root hairs (Favery et al. 2001; Galway et al. 2011; Kim et al. 2007; Park et al. 2011; Wang et al. 2001; Li et al. 2016; Bernal et al. 2008). These tubular cells direct secretion of flexible cell wall materials to their tips, allowing for a process of localized cell expansion known as polarized tip growth (Rounds and Bezanilla 2013; Gu and Nielsen 2013).

*AtCSLD5* and its orthologs are required for normal growth of stems and leaves (Bernal et al. 2007; Hunter et al. 2012; Luan et al. 2011; Yoshikawa et al. 2013; Wu et al. 2010; Hu et al. 2010; Yang et al. 2016; Li et al. 2009). Recently it was discovered that *atcsld5* mutants have primary defects in cytokinesis (Yang et al. 2016; Gu et al. 2016; Hunter et al. 2012) and that the gene is regulated by the cell cycle (Gu et al. 2016; Yoshikawa et al. 2013). In root hairs of Arabidopsis, CSLDs are tip localized, whereas CESAs localize to subapical regions (Park et al. 2011). In cytokinesis, AtCSLD5 accumulates earlier in cell plate development than CESAs, suggesting that it may be responsible for depositing a scaffold upon which CESAs deposit cellulose microfibrils (Gu et al. 2016). Root hair tip growth and AtCSLD3 localization are not affected by isoxaben (Park et al. 2011), which specifically targets CESA proteins (Scheible et al. 2001). However, tips rupture in the presence of DCB (Favery et al. 2001; Park et al. 2011) and CGA 325′615, a CBI that alters the subcellular localization of both CESAs and CSLD3 in Arabidopsis (Park et al. 2011).

The data described above indicate that CSLDs synthesize cellulose or a cellulose-like polymer required to maintain the integrity of growing root hairs and pollen tubes (Park et al. 2011) and developing cell plates (Gu et al. 2016) through a process that is distinct from CESA-mediated cellulose biosynthesis. It has been suggested that CSLDs differ from CESAs in their ability to interact with microtubules (Yang et al. 2020). Another possibility is that CSLDs synthesize cellulose with distinct properties. Possibilities include non-microfibrillar β-1,4-glucan synthesized by isolated CSLDs or microfibrils with distinct lateral dimensions synthesized by CSCs that differ from the rosette-type with 18 subunits. CSLD proteins isolated from *in vitro* reactions were imaged as particles that resembled one lobe of a rosette CSC, suggesting that CSLDs form a type of CSC, however no fibrillar products were detected (Yang et al. 2020).

*Physcomitrium* (formerly *Physcomitrella*) *patens* is typical of many mosses with a dominant haploid phase consisting of tip-growing protonemal filaments and leafy gametophores composed of cells that expand by diffuse growth (Schumaker and Dietrich 1998). Although CESA activity is required for gametophore development (Goss et al. 2012; Scavuzzo-Duggan et al. 2015), protonemal tip growth is insensitive to the CESA-specific CBI isoxaben, but sensitive to DCB (Tran et al. 2018), which affects both CESA and CSLD proteins (Park et al. 2011). Thus, cellulose detected at the tips of growing protonemal filaments (Berry et al. 2016) may not be synthesized by CESAs. Advantages of *P. patens* as an experimental organism include CRISPR-Cas9 based methods for rapid in-locus tagging of proteins (Mallett et al. 2019; Lopez-Obando et al. 2016) and the ability to capture the continuous development of protonemata and gametophores at exquisite temporal and spatial resolution by live-cell imaging (Wu and Bezanilla 2018; Bascom et al. 2016; Bascom et al. 2018).

It is not known whether CSLDs produce microfibrillar cellulose. However, by analogy with CESAs, this would require that CSLDs associate to form CSCs and move in the plasma membrane. Here we demonstrate the role of *P. patens CSLD6* in cytokinesis and protonemal tip growth, and we show that CSLD6 moves in linear trajectories along the plasma membrane, suggesting that it forms complexes and synthesizes microfibrillar cellulose. Differences between CSLD6 and CESA10 in the rate, duration, and CBI sensitivity of these linear movements indicate that CSLD6 and CESA10 function within distinct structures.

## Results

### Moss CSLDs diversified independently from seed plants and have distinct expression patterns

A phylogenetic tree was constructed to examine the diversification of CSLDs in *P. patens* (Roberts and Bushoven 2007); selected non-flowering plant species with sequenced genomes including the lycophyte *Selaginella moellendorffii* (Harholt et al. 2012), the liverwort *Marchantia polymorpha* (CSLD sequences identified by BLAST) and the conifer *Picea abies* (Yin et al. 2014); and angiosperms in which CSLDs have been functionally characterized including Arabidopsis (Richmond and Somerville 2000), cotton (Li et al. 2017), *Populus* species, rice and maize (Yin et al. 2014). The tree was rooted with CSLD sequences from the charophyte green alga *Coleochaete orbicularis* (Mikkelsen et al. 2014). The tree (Figure S1) reveals that the CLSD family diversified independently in mosses, lycophytes, liverworts and seed plants. As shown previously (Hunter et al. 2012), seed plant CSLDs cluster in three clades that correspond to distinct mutant phenotypes related to pollen tube development (Doblin et al. 2001; Bernal et al. 2008; Wang et al. 2011; Moon et al. 2018), root hair development (Favery et al. 2001; Galway et al. 2011; Kim et al. 2007; Park et al. 2011; Wang et al. 2001; Li et al. 2016; Bernal et al. 2008; Peng et al. 2019), and general growth effects, in some cases attributed to defects in cytokinesis (Bernal et al. 2007; Hunter et al. 2012; Luan et al. 2011; Yoshikawa et al. 2013; Wu et al. 2010; Hu et al. 2010; Yang et al. 2016; Gu et al. 2016; Samuga and Joshi 2004). The divergence of *P. patens* CSLDs into two clades, one containing CSLD2 and CSLD6 and the other containing the six remaining *P. patens* CSLDs (Figure S1), is supported by synteny analysis (Figure S2). Analysis of *P. patens* microarray data available in PEATmoss (Fernandez-Pozo et al. 2020) shows distinct expression patterns for members of these clades with *CSLD2* and *CSLD6* enriched in leafy gametophores and the others enriched in filamentous protonemata (Figure S3). Overall, mRNA expression was somewhat higher for *CSLD2* than for *CSLD6*. RNAseq data in PEATmoss confirmed gametophore enrichment for *CSLD2*, but *CSLD6* transcripts were not detected.

### CSLD2 and CSLD6 are redundantly required for gametophore cellular patterning

To investigate functional specialization within the *P. patens* CSLD family, we used homologous recombination to knock out *CSLD2* (Pp3c25_12650V1.1) and *CSLD6* (Pp3c6_4060V1.1), close paralogs that are more highly expressed in gametophores. Multiple independent *csld2*KO and *csld6*KO lines (Figure S4) produced protonemal colonies and gametophores with normal morphologies (Figure 1A-F; Figure S5). In contrast, the gametophores of double *csld*2/6KOs had aberrant phyllids that were smaller than wild type with multiple defects related to cell expansion and cell adhesion (Figure 1G, H; Figure S5). The mildest of these defects were “rosette-like” structures where cells elongated in a radial pattern centered on small cell separations (Figure 1H, J), instead of parallel to the phyllid axis (Figure 1I). These rosette-like structures were also observed occasionally in *csdl2*KO phyllids that were otherwise normal in appearance (Figure 1F). More severe defects observed only in *csld*2/6KOs, including tubular protrusions that appeared to form from continued expansion and division of the radially elongated cells surrounding small cell separations (Figure 1H, K; Figure S6), larger cell separations (Figure 1L), truncated midribs (Figure 1M), and formation of protonema-like filaments along the phyllid margins (Figure 1N). No incomplete cell walls were detected. Normal leaf morphology was restored when a *csld2/6*KO line was transformed with a vector driving expression of either *CSLD2* or *CSLD6* with the native promoter (Figure S7).

**Figure 1:**
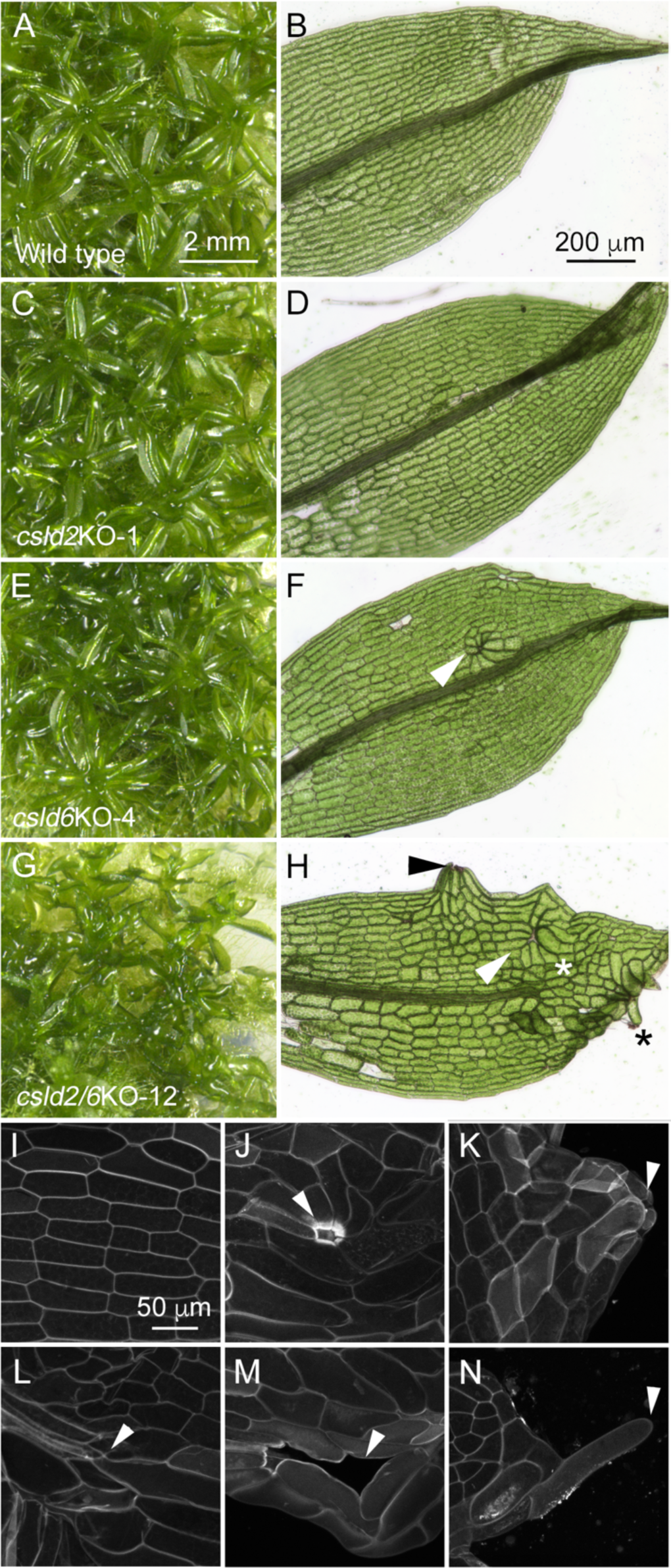
*CSLD2* and *CSLD6* are redundant and required for normal phyllid development. Phyllid development proceeds normally in wild type (A, B), *csld2*KO (C, D) and *csld6*KO (E, F) plants. In contrast, the phyllids of double *csld2/6*KO plants (G, H) show a variety of morphological defects including midribs that do not extend to the tips (white asterisk), formation of protonema-like filaments on the leaf margins (black asterisk) and bulges that sometimes extend to form tube-like structures (black arrowhead). Minor defects consisting of cell separations surrounded by cells with altered growth orientation (white arrowheads) are found in *csld6*KO (F) and *csld2/6*KO (H) phyllids. (I-N) Phyllids stained with Pontamine Fast Scarlet 4B (S4B) and imaged with confocal scanning laser microscopy. Cells elongate parallel to the phyllid axis in wild type (I). Cell adhesion and expansion defects in *csld2/6*KO plants (J-N) include “rosette-like” structures that form where cells surrounding a small cell separation elongate in a radial pattern (J, arrowhead), bulges that form as cells surrounding a small separation elongate and divide (K, arrowhead), midribs that end abruptly instead of extending to the leaf tip (L, arrowhead), large cell separations (M, arrowhead) and marginal cells that extend as filaments that superficially resemble protonemata (M, arrowhead).

We also examined protonemal growth in *csld2/6*KOs by morphometric analysis of colonies grown from protoplasts (Vidali et al. 2007; Li et al. 2019). In two independent experiments, we found no significant differences in colony area between wild type and any of the four *csld2/6*KO lines tested (Figure S8).

### CSLD6 localizes to cytoplasmic puncta and the developing cell plate

To observe CSLD localization and behavior in living cells, we tagged the *CSLD6* locus by inserting sequences coding for fluorescent proteins immediately upstream of the start codon (Figure S9). We generated lines with mEGFP (Vidali et al. 2009b) or the rapidly maturing form of mScarlet, mScarlet-I (Bindels et al. 2017) in a variety of strain backgrounds (Table S1). To test whether the N-terminal fusion resulted in a functional protein, we used CRISPR-Cas9 to disrupt *CSLD2* in the mScarlet-CSLD6/mEGFP-tubulin line. Given that *CSLD2* and *6* are redundant, if mScarlet-CSLD6 is functional, we would expect to recover *csld2*KO plants with no observable phenotype changes. Indeed, we found that both protonemal growth and gametophore morphology were indistinguishable from control plants for the two csld2KO alleles we recovered (Figure S10). As expected, mScarlet-CSLD6 expressed robustly in gametophores where it localized to cytosolic punctae and to the developing cell plate (Figure 2A, Movie S1). During interphase, mScarlet-CSLD6 was relatively evenly distributed in the cytosol. However, during cell division, the cytosolic punctae diminished in number and intensity with a concomitant increase in the amount in the developing cell plate (Figure 2A, insets, Movie S1).

**Figure 2.**
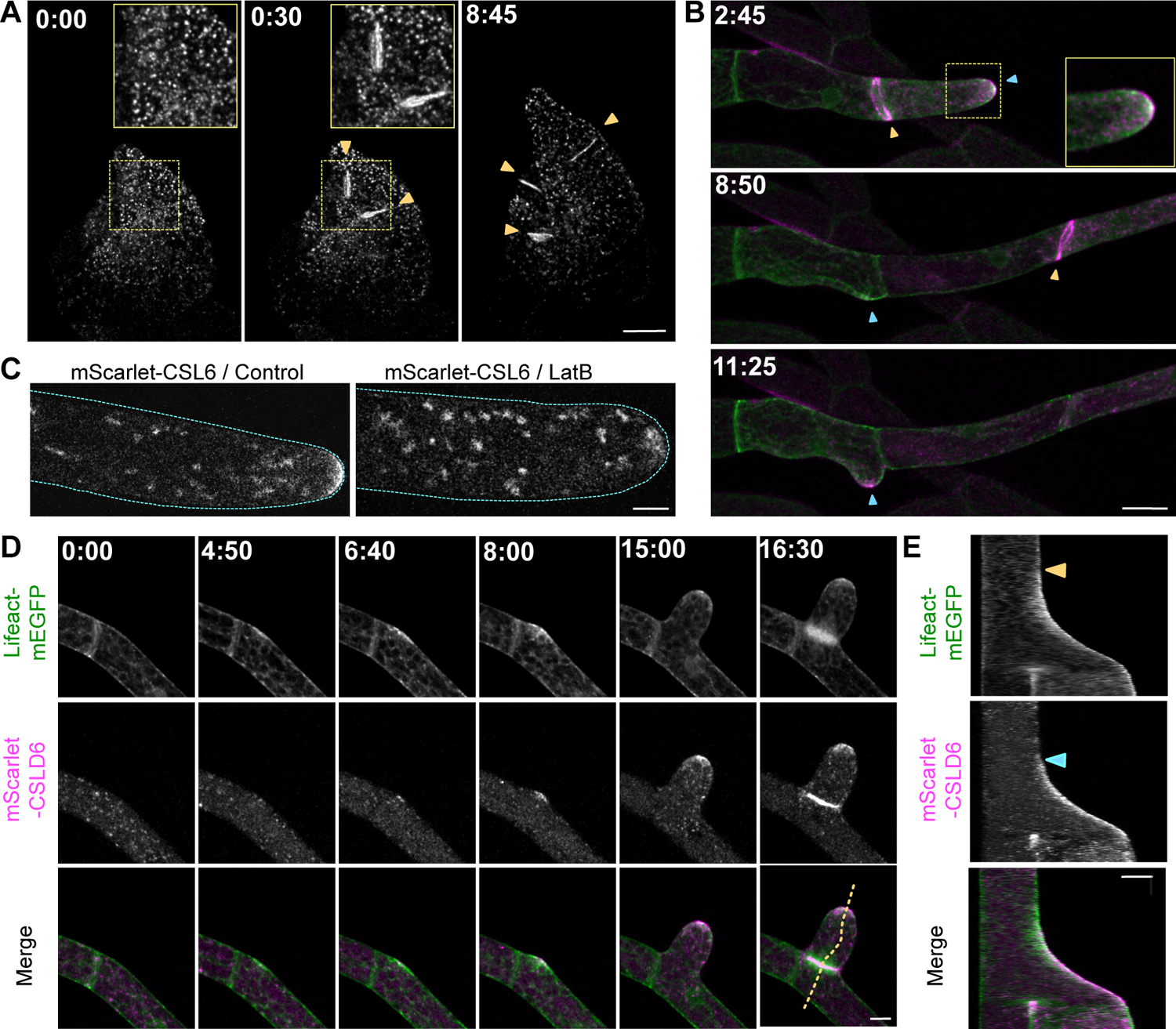
CSLD6 localizes to developing cell plates. At the apex of tip cells CSLD6’s localization is dependent on and follows the accumulation of actin. CSLD6 is tagged with mScarlet on the N-terminus at its endogenous locus. (A) In moss gametophores, CLSD6 is enriched in punctate structures and in the cell plates during cell division (yellow arrow heads.) Insets from the boxed regions reveal the presence of fewer CSLD6 puncta in dividing cells. Scale bar, 20 µm. Time stamps, hour:minute. Also see Movie S1. (B) In moss protonemata, mScarlet-CSLD6 (magenta) accumulates at the cell apex (blue arrow heads) and at the site of cell division (yellow arrow heads.) Actin is labeled with lifeact-mEGFP (green), which also accumulates near the cell apex and at the site of cell division. Inset from the boxed region reveals that CSLD6 labels the plasma membrane, while actin is in the cytosol at the cell apex. Images are maximum projection of z-stacks from a time-lapse acquisition. Inset is from the medial plane. Scale bar, 20 µm. Time stamps, hour:minute. Also see Movie S2 (C) mScarlet-CSLD6 in control and LatB-treated protonemal apical cells. Cyan dotted line outlines the cells. Scale bar, 5 µm. Images are from the medial focal plane. (D) During branch formation, actin appeared (4:50) before CSLD6 accumulation (6:40). Cell expansion occurs (8:00) after actin and CSLD6 accumulation. Images are maximum projections of z-stacks from a time-lapse acquisition. Scale bar, 10 µm. Time stamps, hour:minute. Also see Movie S3. (E) Kymographs are generated along the yellow dash line in (C). In the kymographs, actin appears at the cell apex (yellow arrow heads) before CLSD6 (blue arrow heads) and cell expansion. Scale bars, 10 µm (horizontal) and 2 hours (vertical).

Surprisingly, based on absence of an altered protonemal phenotype in *csld2/6*KO mutants, and in contrast to published *CSLD6* transcript levels available in PEATmoss (Fernandez-Pozo et al. 2020), mScarlet-CSLD6 was well-expressed in protonemata where it also localized to cytosolic punctae. In actively growing protonemal tip cells, mScarlet-CSLD6 accumulated at the apical plasma membrane and along the developing cell plate (Figure 2B, Movie S2), as did mEGFP-CSLD6 (Figure S11). In tip cells, mScarlet-CSLD6 punctae were more concentrated towards the cell apex. However, the level of mScarlet-CSLD6 increased in subapical cells and it accumulated in emerging branches (Figure 2B, Movie S2). Actin, which is essential for polarized growth, accumulates along with secretory vesicles at the apex of tip growing cells, just below the plasma membrane (Vidali et al. 2009a; Bibeau et al. 2020). Imaging of actin, labeled with lifeact-mEGFP, and mScarlet-CSLD6 revealed that CSLD6 punctae accumulate with actin at the cell tip. A sub-population of mScarlet-CSLD6 did not correlate with actin and was found to concentrate on the apical plasma membrane (Figure 2B, inset, Movie S2). However, this membrane population disappeared when cells were treated with latrunculin B (LatB), a drug that depolymerizes the actin cytoskeleton (Figure 2C). These data suggest that actin is required for proper delivery of CSLD to the plasma membrane.

Branch initiation is an excellent model to study the molecular requirements for establishing a new site of polarized growth. Using time-lapse confocal imaging we found that lifeact-mEGFP localizes to the presumptive branch two hours before there is any evidence of cell expansion at that site (Figure 2D, E, Movie S3), similar to previous observations that actin localizes to new polarized sites hours before expansion occurs (Wu and Bezanilla 2018). Notably, mScarlet-CSLD6 was recruited to the same site coincident with emergence of the branch (Figure 2D, E, Movie S3), suggesting that CSLD activity alters cell wall properties in a way that enables expansion.

### CSLD6 movement along linear trajectories in the plasma membrane is sensitive to DCB

We used variable angle epifluorescence microscopy (VAEM) to image CSLD6 behavior at the plasma membrane. For VAEM, we imaged mEGFP-CSLD6 to directly compare it to CESA10 (Pp3c9_2670V1.1), a CESA that is well-expressed in protonemata (Tran and Roberts 2016a). We tagged the *CESA10* locus by inserting mEGF immediately upstream of the start codon (Figure S12). Similar to CESA5 and CESA8 (Tran et al. 2018), we found that mEGFP-CESA10 localized to discrete dots on the plasma membrane that move in linear trajectories (Figure 3A, Movie S4). mEGFP-CSLD6 also localized to discrete dots on the plasma membrane (Figure 3A, Movie S4). However, in contrast to CESA10, CSLD6 movements were significantly faster and shorter in duration (Figure 3A-C, Movie S4). Quantitative analysis of CSLD6 and CESA10 particle motility by particle tracking revealed that, on average, CSLD6 particles had a wider distribution of velocities and moved significantly faster than CESA10 particles (Figure 3B).

**Figure 3.**
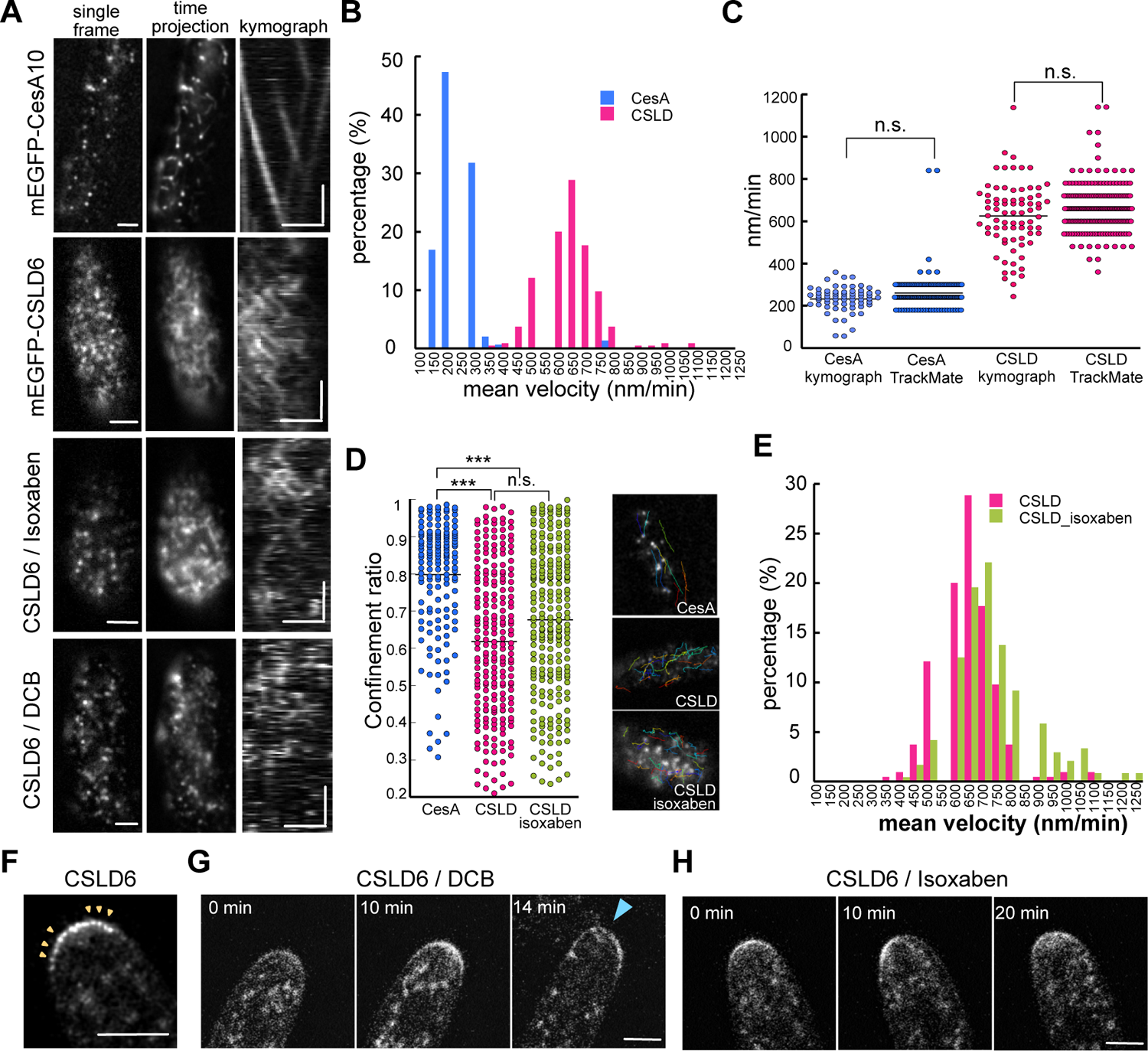
CSLD6 moves in linear trajectories on the plasma membrane, which are specifically inhibited by DCB. (A) Moss protonemata expressing mEGFP-CESA10 or mEGFP-CSLD6 imaged with VAEM. 20µM isoxaben did not affect CSLD6 particle movement, but treatment with 10 µM DCB treatment inhibited CSLD6 particle motility. Scale bars, 2 µm (horizontal) and 2 min (vertical). Kymographs were generated along a trajectory in the time projection. Also see Movies S4 and S5. (B) Histogram of particle velocity as determined by particle tracking with the Fiji plugin TrackMate. On average CSLD6 particles moved faster than CESA10. (C) Velocity measurements from kymographs or from particle tracking are the same. n.s. denotes no significant difference as determined by a student-t test (D) Confinement ratio shows that CESA10 trajectories are straighter than CSLD6 trajectories. Asterisks denote p<0.001as determined by a one-way ANOVA with a Tukey *post hoc* test (α=0.05) and n.s. denotes no significant difference. (E) Histogram of CSLD6 particle velocity with or without isoxaben treatment as determined by particle tracking with the Fiji plugin TrackMate. (F-H) Single focal plane confocal time-lapse image of protonemata expressing mScarlet-CSLD6 with no drug (F), 20µM DCB (G) or 20µM isoxaben (H). Also see Movie S6. (F) CSLD6 accumulates at the cell apex and appeared as puncta (yellow arrow heads). (G) With DCB treatment, CSLD6 still accumulates at the cell apex but the tip cell ruptures (blue arrowhead.) (H) Isoxaben treatment does not cause cell rupture and does not affect CSLD6 localization to the cell tip.

Notably, particle tracking velocity measurements were statistically indistinguishable from kymograph measurements (Figure 3C). Using the particle tracking data, we quantified the confinement ratio, which is a ratio of the actual trajectory distance divided by the straight-line distance between the start and end point of the trajectory. The closer the ratio is to 1, the straighter the trajectory. We found that CSLD6 trajectories were significantly more circuitous than CESA10 (Figure 3D). These data indicate important facets of CSLD6 activity. First, they show that CSLD6 translocates within the plasma membrane. Second, CESA10 and CSLD6 trajectories were separable in speed, duration, and pathway, suggesting that these proteins do not co-assemble.

CBIs such as DCB and isoxaben, have been used to inhibit CESAs *in planta* (Tateno et al. 2016). In *P. patens* protonemata, treatment with DCB or isoxaben substantially reduced CESA linear movements (Tran et al. 2018). We tested whether CSLD6 movements are also sensitive to DCB and isoxaben. VAEM imaging revealed that DCB treatment inhibited mEGFP-CSLD6 linear trajectories, but isoxaben treatment had no effect (Figure 3A, D, E, Movies S5). In addition to not affecting velocity of CSLD6 particles, isoxaben also did not alter the confinement ratio (Figure 3D). This differential sensitivity to DCB and isoxaben provides a tool for distinguishing the effects of CSLD and CESA inhibition. In tip growing cells of Arabidopsis and moss, CSLD6 accumulates at the apical plasma membrane (Figure 3F; Park et al. 2011) and tip rupture is induced by DCB, but not by isoxaben (Figure 3G; Favery et al. 2001; Park et al. 2011; Tran et al. 2018; Rudolph et al. 1989). Imaging cells immediately after exposure to DCB revealed that cells accumulated CSLD6 at the apical plasma membrane before rupture (Figure 3G, Movie S6). In contrast, isoxaben treatment did not affect CLSD6 accumulation at the tip (Figure 3H, Movie S6) and did not affect growth or lead to cell rupture. Together these data suggest that distinct CSLD and CESA membrane complexes are differentially sensitive to CBIs and that CSLDs are required for cell wall integrity at the cell apex.

### CSLDs strengthen the developing cell plate

To probe CSLD function during cytokinesis, we used cells that expressed mScarlet-CSLD6 along with a microtubule marker (mEGFP-tubulin) and stained for callose (using aniline blue) as a cell plate marker. We imaged cytokinesis in protonemata to take advantage of superior temporal and spatial resolution that can be achieved in these cells and used DCB treatment to examine the effects of acute inhibition of CSLD activity. During cell division, CSLD6 began accumulating in the midzone of early phragmoplasts. Fully expanded phragmoplasts exhibited the maximal CSLD6 signal. As the aniline blue signal increased, we observed a concomitant decrease in CSLD6, showing that it functions in early stages of cytokinesis before callose accumulates (Figure 4A, Movie S7). Although DCB had no effect on the timing of mScarlet-CSLD6 accumulation during phragmoplast expansion, we observed that fully expanded phragmoplasts were deformed, often bending in the middle, but subsequently flattened out as callose accumulated (Figure 4B, Movie S8). Like inhibition of CSLD movement (Figure 3A) and tip rupture (Tran et al. 2018), phragmoplast bending was observed only with DCB treatment and not with isoxaben treatment (Figure S13). Together, these results suggest that CSLD activity contributes to the rigidity of the nascent cell plate during phragmoplast expansion and that any role of CESAs is minor in comparison.

**Figure 4.**
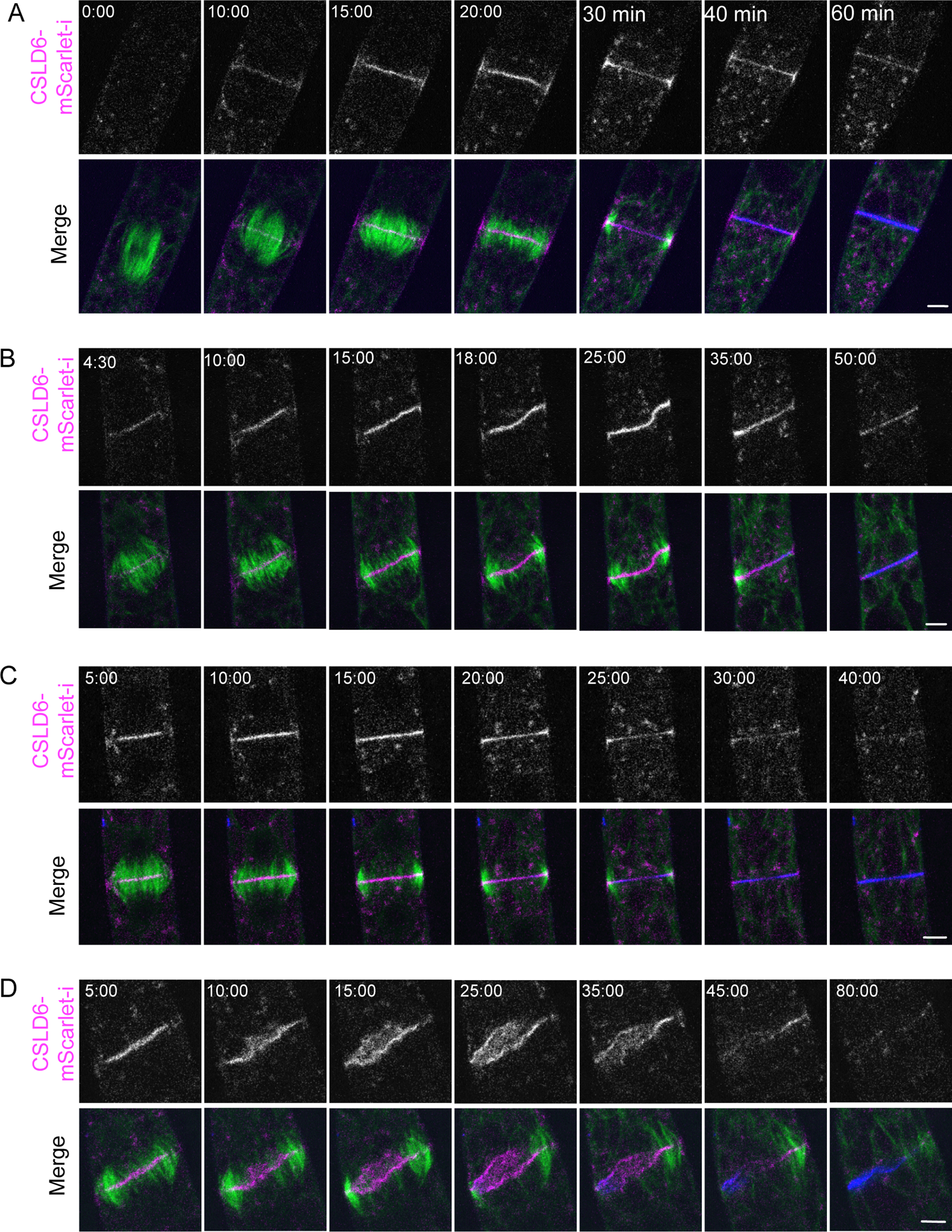
CSLD activity and actin stabilize the nascent cell plate. Cell divisions in moss protonemata expressing mScarlet-CSLD6 (magenta in merge) and mEGFP-tubulin (green in merge) and stained for callose with aniline blue (blue in merge). (A) Cell without drug treatment. Also see Movie S7. (B) Cell treated with 10µM DCB. The cell plate buckles (18:00 and 25:00) but straightens again afterward (35:00 and 50:00). Also see Movie S8. (C) Cell treated with 25 µM LatB. Also see Movie S9. (D) Cell treated with 25 µM LatB and 10 µM DCB. Also see Movie S10. Scale bars, 5 µm. Time stamps, min:sec.

Both microtubules and actin filaments have been hypothesized to direct vesicle trafficking as well as influencing cell plate positioning and structural stabilization (Smertenko et al. 2017). This proposed role in reinforcing the nascent cell plate, prompted us to test whether actin contributed to straightening the bent cell plates in DCB-treated cells. Depolymerizing actin with LatB had no effect on the timing of CSLD6 accumulation at the cell plate (Figure 4C, Movie S9). We also observed that mScarlet-CSLD6 accumulates in the phragmoplast midzone before lifeact-mEGFP (Figure S14). Thus, in contrast to the cell tip, CSLD6 localization to the nascent cell plate does not depend on actin. Division proceeded relatively normally in LatB-treated apical cells (Figure 4C, Movie S9). While difficult to represent in still images, we did notice that the CSLD6 signal appeared wavier during cell plate maturation in LatB-treated cells (Movie S9), suggesting that the nascent cell plate might be slightly less rigid without actin.

Notably, treatment with both LatB and DCB caused more substantial cell plate destabilization than either LatB or DCB treatment alone (Figures 4B-D, Movie S8-S10). LatB did not affect the timing of DCB-induced deformation of cell plates labeled with mScarlet-CSLD6, which still began when the phragmoplast microtubules reached the cell cortex (Figure 4B, D). However, without actin, the cell plates did not flatten out and continued to buckle and twist, ultimately appearing to partially fragment (Figure 4D, Movie S10, Figure S11). Depolymerization of phragmoplast microtubules was delayed. Callose staining occurred later than in controls and was uneven (Figure 4D, Movie S10, Figure S15). In contrast to DCB, isoxaben did not enhance LatB destabilization of cell plates (Figure S13), further supporting that CESA activity does not contribute to strengthening the nascent cell plate. Dramatic defects in cell plate formation induced by acutely inhibiting CSLD and actin suggest that CSLD and actin redundantly function to stabilize the nascent cell plate during cell plate expansion.

## Discussion

### CSLD movement in the plasma membrane is consistent with complex formation and microfibril synthesis

Earlier suggestions that CSLDs form complexes and synthesize microfibrillar cellulose were based on several types of indirect evidence. Cellulose is required to maintain tip integrity in root hairs, and growing evidence indicates that this cellulose is synthesized by CSLDs (Park et al. 2011; Gu and Nielsen 2013; Galway et al. 2011; Yang et al. 2020). Cellulose deposited at the tips of root hairs is fibrillar, yet structurally distinct from the cellulose deposited along the root hair shank (Mulder et al. 2004) and root hair tip growth was disrupted when assembly of these microfibrils was inhibited by recombinant cellulose binding domains (Shpigel et al. 1998).

Mutating one of the two CSLDs required for root hair tip integrity resulted in a patchy distribution of cellulose suggestive of disruption of microfibril assembly (Galway et al. 2011), possibly because CSLD complex formation was blocked. More recently, particles resembling the lobes of rosette CSCs were isolated from *in vitro* CSLD reactions, but fibrils were not detected (Yang et al. 2020). CSLD complex formation is also suggested by intragenic complementation of the *csld1* mutant phenotype in *Lotus japonicus* (Karas et al. 2021). The possibility that CSLDs move in the plasma membrane was suggested based on the premise that DCB-induced expansion of the tip-localized AtCSLD3 zone in root hairs is attributable to inhibition of tip-directed CSLD movement (Park et al. 2011). VAEM imaging of GFP-CSLD6 in *P. patens* protonemata (Figure 3, Movies S4-5) provides the first direct evidence that CSLDs move in linear trajectories along the plasma membrane. Based on analogy with CESAs (Diotallevi and Mulder 2007) we postulate that this translocation results from CSLD6 enzymatic activity and the production of microfibrils that upon incorporation into the cell wall, pushes the CSLD6 in the plane of the membrane. A structurally distinct CSC formed by CSLDs instead of CESAs could synthesize microfibrillar cellulose with distinct properties that facilitate tip growth and cytokinesis.

### CESA and CSLD are in different complexes

Chimeric proteins consisting of AtCSLD3 with its catalytic domain replaced by the AtCESA6 catalytic domain can rescue the *atcsld3* root hair growth phenotype (Park et al. 2011).

Conversely, a chimera of AtCESA6 with the catalytic domain AtCSLD3 can rescue the *atcesa6* null phenotype. The fluorescently-tagged chimeric protein moved in the plasma membrane as part of a complex that also contained AtCESA3. Although these data indicate that regions outside the catalytic domain of these proteins are responsible for their distinct functions, including complex formation, fibrillar structure of the product, and subcellular targeting (Yang et al. 2020), they do not rule out the possibility that CESAs and CSLDs form hetero-complexes.

We show that CSLD6 movements are faster and shorter in duration as compared to CESA10 and are not inhibited by isoxaben (Figure 3, Movies S4-5). These distinct patterns of movement indicate that CSLD6 and CESA10 function within different structures and produce microfibrillar cellulose at distinct rates. This supports previous observations suggesting separation of CESA and CSLD function in time and space. Freeze fracture EM of *P. patens* protonemal filaments has revealed abundant rosette CSCs associated within subapical regions of protonemal tips (Nixon et al. 2016; Roberts et al. 2012), which is consistent with subapical localization of eGFP-CESA6 in Arabidopsis root hairs (Park et al. 2011) and the well-characterized role of CESAs in deposition of root hair secondary cell walls (Emons 1994; Lindeboom et al. 2008). In contrast, CSLDs are localized to the tips of both *P. patens* protonemata (Figure 2, 3) and Arabidopsis root hairs (Park et al. 2011). Similarly, rosettes (Roberts et al. 2012) have been observed and GFP-CESAs are more abundant (Gu et al. 2016; Miart et al. 2014) in the late stages of cell plate maturation, whereas CSLDs are present in the early cell plates in *P. patens* (Figure 2, 4) and Arabidopsis (Gu et al. 2016).

### Role of CSLD6 in tip growth, cell division and cell expansion

In Arabidopsis and other seed plants, *csld* mutations cause defects in either cytokinesis or tip growth. Here we show that *P. patens* CSLD6 is associated with both of these processes. First, mScarlet-CSLD6 is localized to protonemal cell tips and to cell plates of both protonemata and developing phyllids (Figure 2). Second, DCB inhibited mEGFP-CLSD6 movement in the plasma membrane and induced tip rupture (Figure 3, Movies S4-6), and destabilized developing cell plates (Figure 4, Movies S7-10). Notably, CSLD stabilization of the nascent cell plate is synergistic with actin. Acute inhibition of either CSLD or actin did not dramatically alter cytokinesis (Figure 4). However, acute inhibition of both CSLD and actin resulted in cytokinesis failure (Figure 4), suggesting that actin provides cytosolic tension while CSLD provides cell plate rigidity. Isoxaben, which specifically targets CESAs (Scheible et al. 2001), had no effect on any of these processes, suggesting that the effects of DCB are attributable to CLSD inhibition. Collectively, these data support the hypothesis that the product of CSLD activity stabilizes nascent cell walls formed during tip growth and cytokinesis in *P. patens*.

Mutation analysis revealed that CSLD2 and CSLD6 function redundantly to maintain normal phyllid development. Although *csld2*KOs and *csld6*KOs have little or no obvious phenotype, double *csld2/6*KOs produce aberrant phyllids with defects in cell adhesion and patterning. These double mutants did not have incomplete walls like plants resulting from mutations in *AtCSLD5* or its ortholog *ZmCLSD*1, which are associated with cytokinesis in Arabidopsis (Gu et al. 2016) and maize (Hunter et al. 2012). However, *csld2/6*KOs phyllids do display cell separations, which may be an alternative consequence of disrupted cytokinesis. Many of the cell patterning defects appear to result from changes in the direction of expansion of cells directly adjacent to the cell separations (Figure 1). The separations almost certainly alter the stress-strain relationships between adjacent cells and it is known that plant cells alter their direction of cell expansion in response to mechanical signals (Hamant and Haswell 2017). Thus, the patterning defects observed in phyllids may be an indirect consequence of cell separation, resulting from altered tissue mechanical forces and consequent changes in the polarity of cell expansion.

Mutations that affect *AtCSLD5* or its orthologs in other seed plants also alter cell patterning (Gu et al. 2016; Hunter et al. 2012; Yoshikawa et al. 2013; Yang et al. 2016). Effects include epidermal bulges on maize leaves that superficially resemble the bulges we observed on moss phyllids (Hunter et al. 2012) and enlarged and misshapen bulliform, bundle sheath, and stomatal lineage cells in rice (Yoshikawa et al. 2013). These data suggest that CSLD impairment may impact cell expansion. Interestingly, Yang et al (2016) reported linkage composition changes for all polysaccharide classes in the meristems of Arabidopsis *csld* mutants. Compensatory alteration of cell wall composition is a common response to chemical or genetic inhibition of cellulose synthesis that is thought to be mediated by a cell wall integrity sensing system (Anderson and Kieber 2020). This response may underly the varied developmental defects that result from *csld* mutations in *P. patens* and other plant species.

Although the *csld2/6*KOs had no obvious protonemal phenotypes, localization of CSLD6 to protonemal cell tips and cell plates suggests that *CSLD6* functions redundantly with one or more of the six other *CSLD*s that are expressed in protonemata. Live cell imaging experiments employing lines expressing lifeact-mEGFP along with mScarlet-CSLD showed that actin accumulates 1-2 hours before CSLD at new growth sites. Notably, cell expansion did not visibly occur until CSLD6 was recruited to the growth site, suggesting that CSLD6 activity is in part required to produce extensible cell walls. Furthermore, actin depolymerization with LatB demonstrated that during tip growth actin maintains CSLD6 accumulation at cell tips, similar to what has been shown previously in Arabidopsis root hairs (Park et al. 2011). In contrast during cell division, CSLD6 accumulates in the phragmoplast before actin and in the presence of LatB, suggesting that actin-independent trafficking recruits CSLD6 to the developing cell plate. During cell division, actin instead provides tension forces that help to strengthen the cell plate during expansion. Our data demonstrate that the same protein is trafficked via actin-dependent and - independent pathways to distinct subcellular sites, the cell tip and the phragmoplast. Ultimately CSLD activity contributes to the balance of cell wall flexibility and strength facilitating expansion of the cell tip and, together with actin, the nascent cell plate.

### CESA/CSLD diversification

Phylogenetic analysis confirms that the *CSLD* families of mosses and seed plants diversified independently from a single ancestral gene. CSLD families from both lineages include members that function in tip growth and cytokinesis, suggesting that both functions are ancestral. In the seed plant lineage, CSLD orthologs from Arabidopsis and rice have similar expression patterns in pollen (Wang et al. 2011; Moon et al. 2018; Bernal et al. 2008), root hairs (Kim et al. 2007; Bernal et al. 2008) or in dividing cells (Gu et al. 2016; Yoshikawa et al. 2013; Yang et al. 2016) indicating that subfunctionalization (*i.e.*, evolution of differential expression patterns resulting from promoter modification) preceded the divergence of monocot and eudicots. In *P. patens* the two CSLD clades are distinguished by preferential expression in gametophores or protonemata (Figure S2, S3) indicating that subfunctionalization preceded the first whole genome duplication. In Arabidopsis, single *csld* mutations have distinctive phenotypes, consistent with limited redundancy (Hunter et al. 2012), although analysis of double and triple mutants showed some overlapping function (Yang et al. 2016; Yin et al. 2011). In *P. patens*, CSLD6 is redundant with CSLD2, and also appears to have functional overlap with one or more of the remaining six CSLDs.

## Materials and methods

### Vector construction

All primer pairs are shown in Table S2, along with annealing temperatures used for PCR. Amplification with Taq Polymerase (New England Biolabs, Ipswich, MA, USA) included a 3 min denaturation at 94°C; 35 cycles of 15 s at 94°C, 30 s at the annealing temperature, and 1 min/kbp at 72°C. Amplification with Phusion Polymerase (New England Biolabs) included a 30 s denaturation at 98°C; 35 cycles of 7 s at 98°C, 7 s at the annealing temperature, and 30 s/kbp at 72°C.

The CSLD2KO and CSLD6KO vectors were constructed using Gateway Multisite Pro cloning (Invitrogen, Grand Island, NY, USA) as described previously (Roberts et al. 2011). Flanking sequences 5’ and 3’ of the coding regions were amplified with appropriate primer pairs (Table S2) using Phusion DNA polymerase (New England Biolabs) and cloned into pDONR 221 P1-P4 and pDONR 221 P3-P2, respectively, using BP Clonase II (Invitrogen). The *hph* selection cassette was amplified from BHNSR (gift of Didier Schaefer, University of Neuchâtel) and cloned into pDONR 221 P3r-P4r. All entry clones were sequence-verified. Entry clones with *CSLD2* and *CSLD6* flanking sequences in pDONR 221 P1-P4 and pDONR 221 P3-P2 were linked with the entry clone containing the *hph* selection cassette or an *nph* selection cassette (Norris et al. 2017), respectively, and inserted into pGEM-gate (Vidali et al. 2009b) using LR Clonase II Plus (Invitrogen). The resulting plasmids were cut with BsrGI for transformation into *P. patens*.

Expression vectors for HA-tagged PpCSLDs under control of their native promoters were constructed using Gateway Multisite Pro cloning (Invitrogen). The *CSLD2* and *CSLD6* coding sequences were amplified from cDNA clones pdp04669 and pdp21814 (RIKEN BioResource Center, Tsukuba, Ibaraki JP), respectively, using Phusion DNA polymerase (New England Biolabs) with forward primers containing a single hemagglutinin (HA) tag and appropriate reverse primers (Table S2), and cloned into pDONR 221 P5-P2 using BP Clonase II (Invitrogen). Similarly, the *CSLD2* and *CSLD6* promoters (approximately 2 Kb upstream of the start codons) were amplified from *P. patens* genomic DNA and cloned into pDONR 221 P1-P5r. All entry clones were sequence verified. Promoter entry clones were linked to entry clones containing their respective *HA-PpCSLD* coding sequences and inserted into pSi3(TH)GW (Tran and Roberts 2016b) using LR Clonase II Plus (Invitrogen). These vectors target the expression cassettes to the intergenic 108 locus, which can be disrupted with no effect on phenotype (Schaefer and Zryd 1997). Rescue vectors were cut with SwaI for transformation into a *P. patens csld2/6*KO lines from which the *hph* resistance cassette had been removed (see below).

CRISPR-mediated homology directed repair was used to insert sequences encoding mEGFP or mScarlet-I in frame with the *CSLD6* gene. To clone the protospacers flanking the first intron (Figure S9), we generated a modified version of pMH-Cas9-gate (Mallett et al. 2019). In this modified vector, the Gateway cassette was replaced with the PpU6 promoter followed by two unique restriction sites (PmeI and SnaBI) and the *S. pyogenes* scaffold RNA sequence from pENTR-PpU6p-sgRNA-L1L2 (Mallett et al. 2019), generating pMH-Cas9. Two separate single stranded oligos (Table S2) containing the protospacer sequence and sequences that overlap with pMH-Cas9 were assembled into pMH-Cas9 using Hi-Fi DNA Assembly Mastermix (NEB). The Hi-Fi reaction was transformed into *E. coli* and used directly to inoculate a 50 mL culture. A midi prep isolated a mix of pMH-Cas9-PS1 and pMH-Cas9-PS2 plasmids. This mixture was used to transform moss. The homology plasmids were assembled using HI-Fi DNA Assembly Mastermix (NEB) with four PCR-generated DNA fragments (see Table S2 for primers): ∼800 bp flanking sequences 5’ and 3’ of the start codon, sequences of mEGFP or mScarlet-I, and the pGEM vector. A different vector system was used to insert mEGFP in-frame with the CESA10 gene. The CRISPR-Cas9 vector was constructed as described previously (Mallett et al. 2019). A protospacer targeting a site near the *CESA10* start codon (Table S2) was annealed as described previously (Mallett et al. 2019) and ligated into the pENTR-Ppu6p-sgRNA-L1L2 entry vector using Golden Gate assembly (New England Biolabs, Ipswich, MA) in a 10 μl reaction containing 37.5 ng of entry vector and 1.1 μl of a 1000x dilution of annealed protospacer incubated at 37°C for 1 h and 60°C for 5 min. The entry vector was cloned into pMK-Cas9-gate conferring G-418 resistance using LR Clonase II Plus according to the manufacturer’s instructions (Invitrogen). All plasmids were sequence verified. We used a homology repair plasmid orginally designed for homologous recombination. An entry clone encoding an mEGFP-CESA10 translational fusion in pDONR 221 P5-P2 was constructed by PCR fusion of the *mEGFP* coding sequence amplified from an expression clone containing a pDONR 221 P1-P5r mEGFP entry clone and the *CESA10* coding sequence amplified from expression clone 3X*HA-CESA10* in xt18 using the appropriate primers (Table S2) as described previously (Scavuzzo-Duggan et al. 2015). The entry clone was inserted into pSi3(TH)GW along with the pDONR 221 P1-P5r entry clone containing the *PpCESA10* promoter (Tran and Roberts 2016a).

### Culture and transformation of P. patens

Wild type *P. patens* lines (haploid) derived from the sequenced Gransden strain (Rensing et al. 2008) by selfing and propagation from a single spore in 2011 (GD11) was a gift of Pierre-Francois Perroud, Marburg University. Wild type and transformed *P. patens* lines were cultured on basal medium supplemented with ammonium tartrate (BCDAT) as described previously (Roberts et al. 2011). Protoplasts were prepared and transformed as described previously (Roberts et al. 2011). Stable transformants were selected with 50 μg mL^-1^ G418 (CSLD6KO vector) or 15 μg mL^-1^ hygromycin (CSLD2KO and complementation vectors). The *hph* selection cassette was removed from *csld2/6*KO as described previously (Norris et al. 2017). CRISPR-Cas9 transformation and selection methods were described previously (Mallett et al. 2019).

### Analysis of knockout lines

For PCR analysis of target integration, target-gene disruption, and selection cassette excision, DNA was extracted as described previously (Roberts et al. 2011) and amplified using primers listed in Table S2. Sanger sequencing was employed to sequence PCR products. Colony morphology was analyzed as described previously (Bibeau and Vidali 2014; Vidali et al. 2007; Li et al. 2019).

### Laser Scanning Confocal and Variable Angle Epifluorescence Microscopy

To image mEGFP/mScarlet-CSLD6 localization and dynamics we employed two sample preparation methods. For long term imaging (>2 hours), we used microfluidic imaging chambers (Bascom et al., 2016). Ground protonemal tissue was gently pipetted into the central part of the device followed by an infusion of Hoagland’s medium. Then the chamber was submerged in Hoagland’s medium (4mM KNO_3_, 2mM KH_2_PO_4_, 1mM Ca(NO_3_)_2_, 89 µM Fe citrate, 300 µM MgSO_4_, 9.93 µM H_3_BO_3_, 220 nM CuSO_4_, 1.966 µM MnCl_2_, 231 nM CoCl_2_, 191 nM ZnSO_4_, 169 nM KI, 103 nM Na_2_MoO_4_), and placed under constant 85 µmol photons m^-2^s light^-1^. For short term imaging (< 2 hours), 5- to 8-day old plants regenerated from protoplasts were placed onto an agar pad in Hoagland’s buffer, covered by a glass cover slip and sealed with VALAP (1:1:1 parts of Vaseline, lanoline, and paraffin).

For confocal microscopy, images were acquired on a Nikon A1R confocal microscope system with either a 0.75. NA 20X air objective (Nikon) (gametophore images) or a 1.49 NA 60X oil immersion objective (Nikon) (protonemata images) at room temperature. Image acquisition was controlled by NIS-Element AR 4.1 software (Nikon). Laser illumination at 405 nm was used to excite aniline blue; 488 nm for mEGFP; 561 nm for mScarlet. Emission filters were 525/50 nm for aniline blue and, mEGFP; 595/50 nm for mScarlet. Sequential acquisition was used whenever imaging aniline blue and mEGFP.

For VAEM, images were acquired with a Nikon Ti-E inverted microscope equipped with a TI-TIRF-PAU illuminator, using a Nikon 1.49 NA 100x oil immersion TIRF objective. mEGFP was illuminated with a 488nm laser, and the emission passed through a 525/50 filter. Images were captured with an Andor 897 EMCCD camera. Image acquisition was controlled by Nikon NIS-Elements software. All data was processed with enhanced contrast (0.1% pixel saturation), subtract background and smoothing in Fiji using default settings.

### Particle tracking

mEGFP-CesA10 and mEGFP-CSLD6 particles were imaged with VAEM every 2s for 10-20 minutes. Images were processed before tracking using subtract background and enhance contrast in Fiji. Particle tracking was performed with the Fiji plugin TrackMate (Tinevez et al. 2017). For CesA10, stacks were reduced by 10, resulting in 20 s time intervals between frames. For CSLD6, stacks were reduced by 6, resulting in 12 s time intervals between frames. LoG detector (estimate blob diameter 0.3 µm) and LAP tracker (frame to frame linking max distance 0.5 µm; Penalties: Quality=1; no gap closing allowed) were selected. The filters on tracks were set as follows: duration >60 s; track displacement > 0.5 µm. Mean velocity and confinement ratio of tracks were obtained and plotted in Figure 3.

### Phylogenetic analysis

For phylogenetic analysis, we include *P. patens CSLD* sequences identified previously (Roberts and Bushoven 2007) along with sequences previously identified in the lycophyte *Selaginella moellendorffii* (Harholt et al. 2012) and the charophyte green alga *Coleochaete orbicularis* (Mikkelsen et al. 2014) We included CSLDs from angiosperm species (Yin et al. 2014) in which CSLD function has been investigated, including *Arabidopsis thaliana*, *Gossypium raimodii* (Li et al. 2017), *Populus trichocarpa*, *Zea mays* and *Oryza sativa*. Sequences were aligned using Clustal/W with BLOSUM cost matrix, gap open cost of 10, and gap extension cost of 0.1.

Selected sequences were realigned as described in results. The alignments were edited to remove gaps and adjacent poorly aligned segments (Baldauf 2003) and rapid bootstrapping (1000 replicates) and search for the best-scoring maximum likelihood tree was carried out using RAxML 7.2.8, GAMMA BLOSOM62 protein model. The trees were exported from Geneious 8.1 and edited with Inkscape (https://inkscape.org/en/).

### Accession numbers

Sequence data from this article can be found in the CoGe/Genbank/Phytozome data libraries under accession numbers listed in Figure S1.

## Acknowledgements

This work was supported by a University of Rhode Island Division of Research and Economic Development Bridge Grant to AWR (construction and characterization of *csld* knockout lines) and as part of The Center for LignoCellulose Structure and Formation, an Energy Frontier Research Center funded by the U.S. Department of Energy, Office of Science, Basic Energy Sciences under Award # DE-SC0001090 (construction and characterization of FP-CESA10). Some material is based upon work conducted at a Rhode Island NSF EPSCoR research facility, the Genomics and Sequencing Center, supported in part by the National Science Foundation EPSCoR Cooperative Agreement #EPS-1004057. Additional support was provided by a grant from the National Science Foundation to MB (MCB-1715785).

## Author Contributions

A.W.R. and M.B. designed the research; S.-Z.W., A.M.C., R.L., A.W.R. and M.B. performed research and analyzed data; A.W.R. and M.B. wrote the paper.

## Supplemental Information

### Supplemental Tables

**Table S1.** Lines used in this study.

| Name | Background | Genetic alteration | Translation |
| --- | --- | --- | --- |
| *csld2*KO | Wild type | locus deleted | NA |
| *csld6*KO | Wild type | locus deleted | NA |
| *csld2/6*KO | *csld2*KO-32 | locus deleted | NA |
| *csld2/6*KO-lox | *csld2/6*KO-16 | *hph* selection cassette excised | NA |
| *csld2/6*KO+CSLD2 | *csld2/6*KO-lox | CSLD2 expression cassette inserted in 108 locus | NA |
| *csld2/6*KO+CSLD6 | *csld2/6*KO-lox | CSLD6 expression cassette inserted in 108 locus | NA |
| *csld2/6*KO+EV | *csld2/6*KO-lox | Empty vector inserted in 108 locus | NA |
| *mEGFP-CSLD6* | Wild type | insertion of mEGFP before ATG | Translational fusion of mEGFP-CSLD6 |
| *mScarlet-CSLD6* | Wild type, mEGFP-tubulin, Lifeact-mEGFP | insertion of mScarlet before ATG | Translational fusion of mScarlet-CSLD6 |
| *csld2*KO-4c | mEGFP-tubulin/<br>mScarlet-CSLD6 | 17 bp deletion | Stop at AA64, aa24-64 are irrelevant |
| *csld2*KO-6c | mEGFP-tubulin/<br>mScarlet-CSLD6 | 10 bp deletion | Stop at AA26, aa22-26 are irrelevant |
| *mEGFP-CESA10* | Wild type | Insertion of mEGFP before ATG | Translational fusion of mEGFP-CESA10 |

**Table S2.**
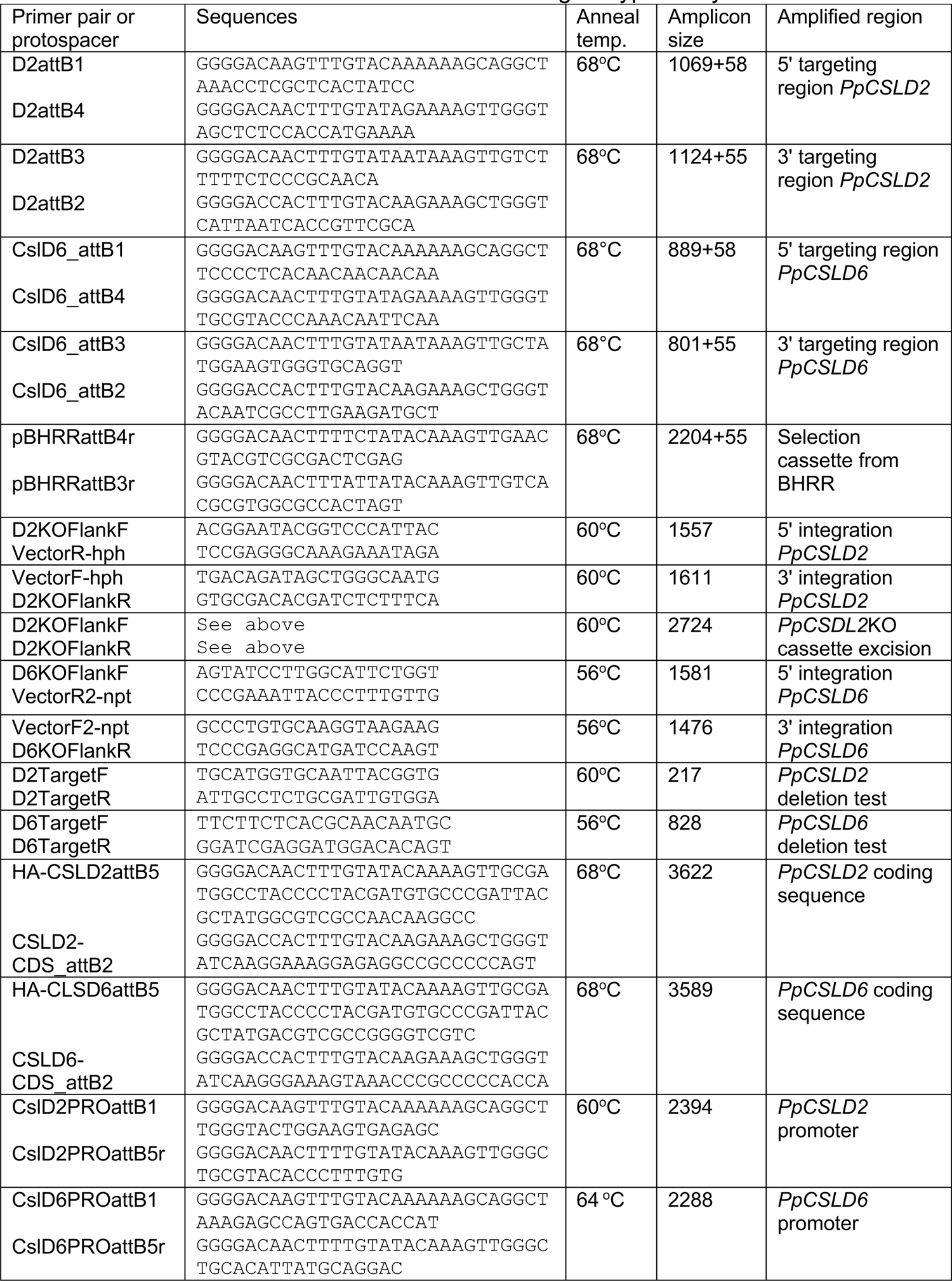
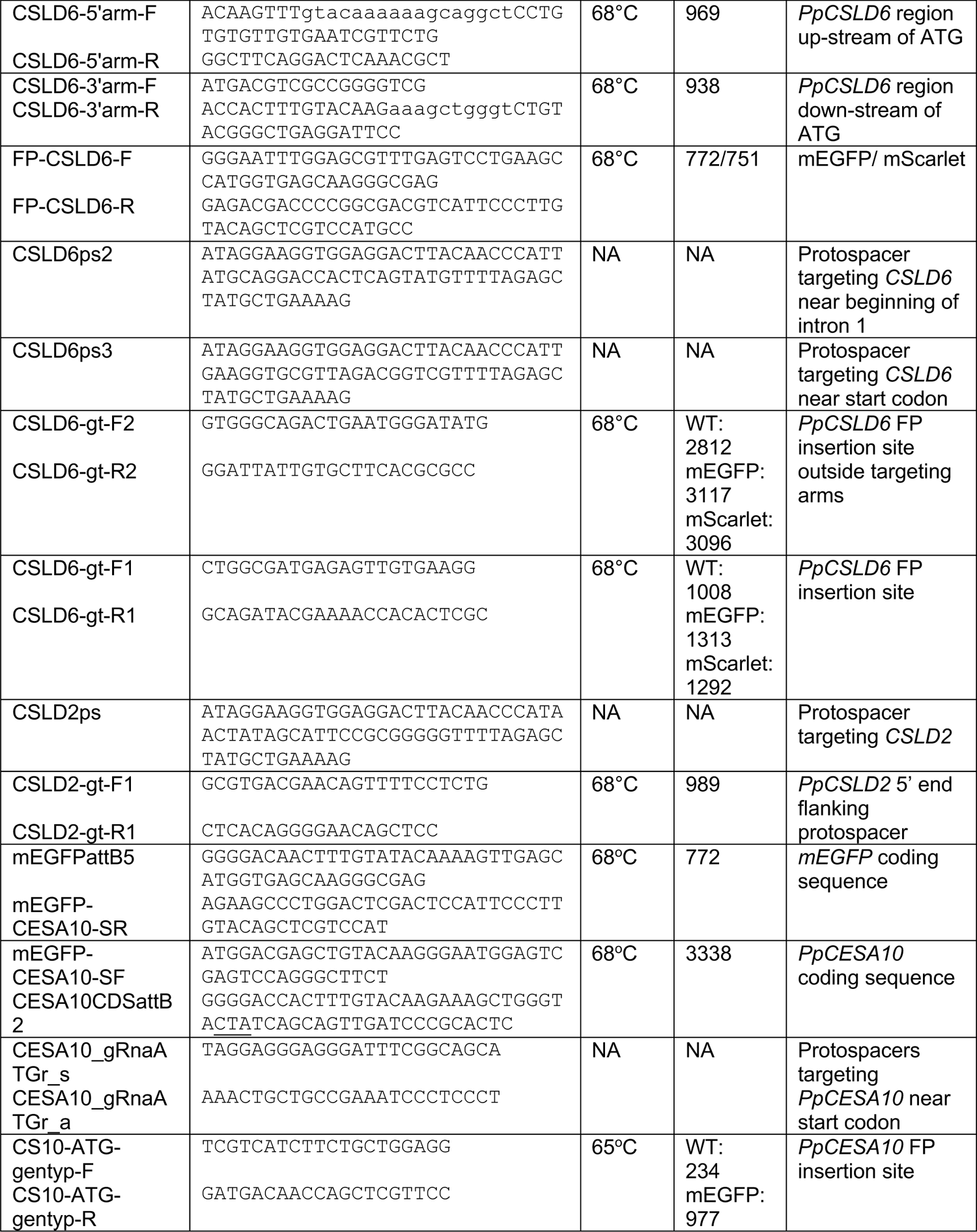
Primers used for vector construction and genotype analysis.

### Supplemental Movie Legends

**Movie S1.** Wild type gametophore expressing mScarlet-CLSD6. Images are maximum projection of z-stacks acquired on a laser scanning confocal microscope. Time interval, 5 minutes. Scale bar, 20 µm. Movie is playing at 8 fps. Also see Figure 2A.

**Movie S2.** Wild type protonemata expressing Lifeact-mEGFP and mScarlet-CLSD6. Images are maximum projection of z-stacks acquired on a laser scanning confocal microscope. Time interval, 5 minutes. Scale bar, 20 µm. Movie is playing at 8 fps. Also see Figure 2B.

**Movie S3.** Wild type protonemata expressing Lifeact-mEGFP and mScarlet-CLSD6. Images are maximum projection of z-stacks acquired on a laser scanning confocal microscope. Time interval, 5 minutes. Scale bar, 20 µm. Movie is playing at 8 fps. Also see Figure 2D.

**Movie S4.** Wild type protonemata expressing GFP-CesA10 and mEGFP-CLSD6. Images are time lapse acquired with VAEM. Time interval, 6 sec. Scale bar, 2 µm. Movie is playing at 15 fps. Also see Figure 3A.

**Movie S5**. Wild type protonemata expressing mEGFP-CLSD6 treated with 10 µM DCB or 20 µM isoxaben. Images are time lapse acquired with VAEM. Time interval, 6 sec. Scale bar, 2 µm. Movie is playing at 15 fps. Also see Figure 3A.

**Movie S6.** Wild type protonemata expressing mScarlet-CLSD6 treated with no drug, 10 µM DCB, or 20 µM isoxaben. The tip cell ruptured with DCB treatment. Images are single focal plane images acquired on a laser scanning confocal microscope. Time interval, 20 sec. Scale bar, 5 µm. Movie is playing at 8 fps. Also see Figure 3F-H.

**Movie S7.** Cell division in tip cell. Wild type protonemata expressing mEGFP-tubulin and mScarlet-CLSD6 were stained with aniline blue. Images are single focal plane images acquired on a laser scanning confocal microscope. Time interval, 1 minutes. Scale bar, 5 µm. Movie is playing at 8 fps. Also see Figure 4A.

**Movie S8.** WT cell treated with 10 µM DCB undergoing cell division. Wild type protonemata expressing mEGFP-tubulin and mScarlet-CLSD6 were stained with aniline blue. Images are single focal plane images acquired on a laser scanning confocal microscope. Time interval, 1 minutes. Scale bar, 5 µm. Movie is playing at 8 fps. Also see Figure 4B.

**Movie S9.** WT cell treated with 25 µM LatB undergoing cell division. Wild type protonemata expressing mEGFP-tubulin and mScarlet-CLSD6 were stained with aniline blue. Images are single focal plane images acquired on a laser scanning confocal microscope. Time interval, 1 minutes. Scale bar, 5 µm. Movie is playing at 8 fps. Also see Figure 4C.

**Movie S10.** WT cell treated with 25 µM LatB and 10 µM DCB undergoing cell division. Wild type protonemata expressing mEGFP-tubulin and mScarlet-CLSD6 were stained with aniline blue. Images are single focal plan images acquired on a laser scanning confocal microscope. Time interval, 1 minutes. Scale bar, 5 µm. Movie is playing at 8 fps. Also see Figure 4D.

### Supplemental Figures

**Figure S1:**
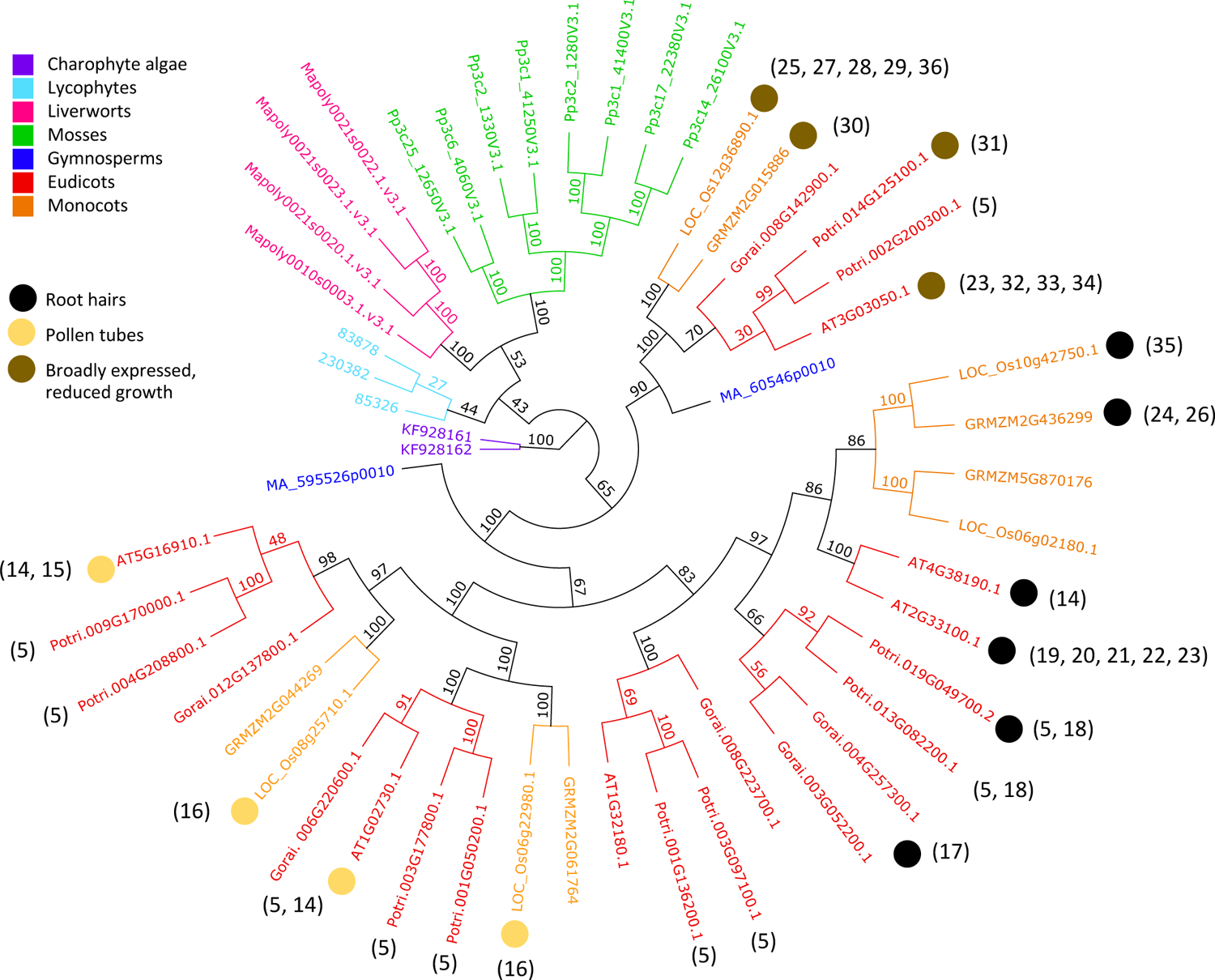
Maximum likelihood tree of CSLD sequences from selected land plant species rooted with green algal CSLD sequences. The CSLD families of mosses, lycophytes, liverworts and charophyte green algae diversified independently. Angiosperm CSLDs group by function (circles) with sequences from other species. Nodes are labelled with bootstrap values (1000 replicates). Species include *Physcomitrium patens*, Pp, CoGe locus IDs^1^; *Coleochaete orbicularis* (GenBank IDs)^2^; *Selaginella moellendorffi* (Phytozome protein IDs)^3^; *Marchantia polymorpha* (Mapoly, Phytozome protein IDs)^4^, *Picea abies* (MA, Phytozome protein IDs)^4^; *Arabidopisis thaliana* (AT, locus IDs)^4^; *Populus trichocarpa* (Potri, Phytozome protein IDs)^5^; *Gossypium raimondii* (Gorai, Phytozome protein IDs)^6^; *Zea mays* (GRMZM, Phytozome protein IDs)^7^ and *Oryza sativa* (Os, locus IDs)^7^.

**Figure S2:**
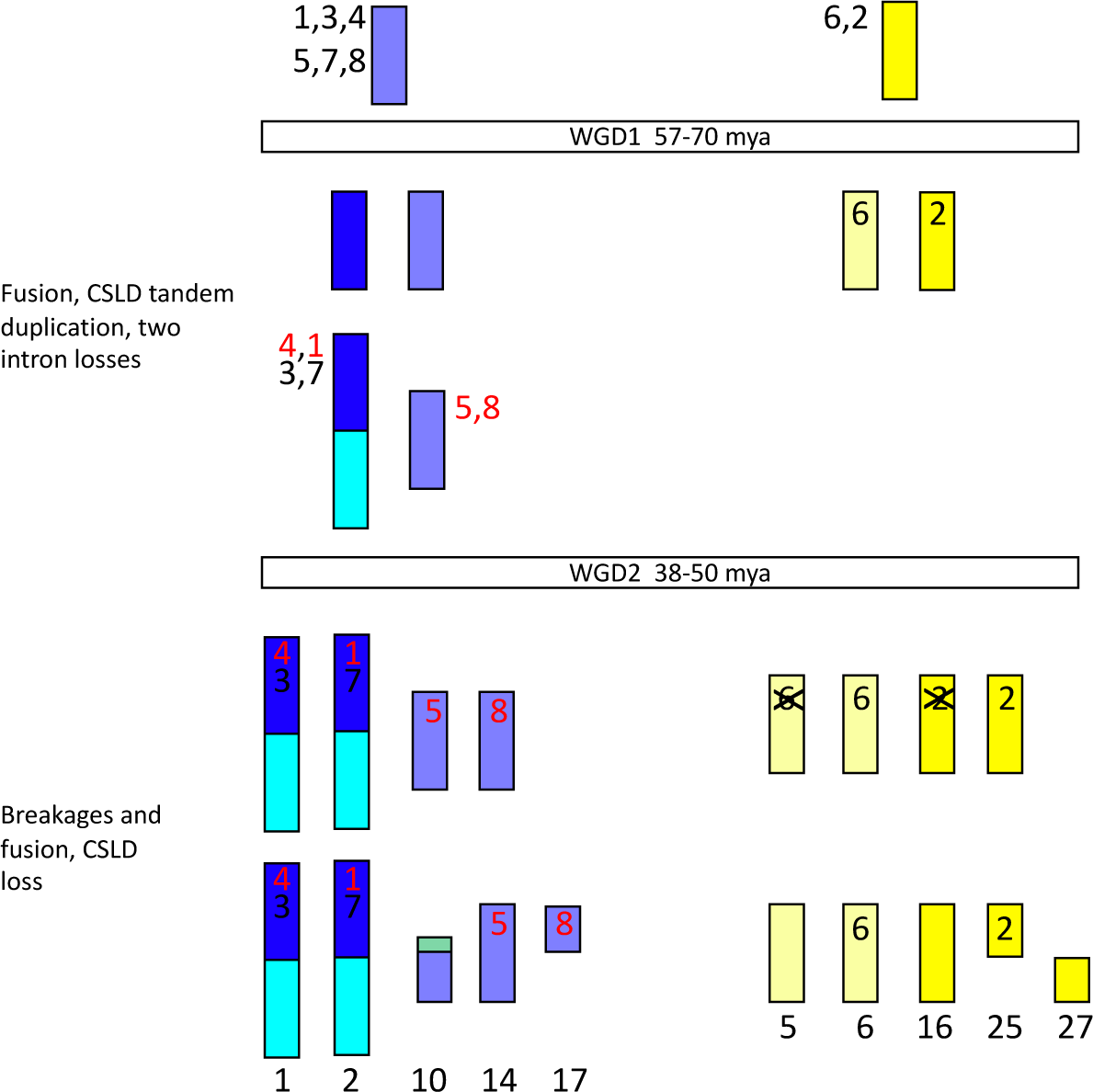
*CSLD2* and *CSLD6* are close paralogs. Synteny analysis of CSLD diversification based on the chromosome-scale assembly of the *P. patens* genome (8). The eight *P. patens* CSLDs reside on chromosomes descended from two of the seven chromosomes proposed to have existed before the first of two whole genome duplications (WGD). *CSLD2* and *6* diverged from a common ancestor in WGD1. Following WGD2 paralogs of *CSLD6* and *CSLD2* were lost from chromosomes 5 and 16, respectively. Duplication of the chromosome carrying the common ancestor of CSD1, 3, 4, 5, 7 and 8 in WGD1 was followed by a fusion affecting the common ancestor of chromosomes 1 and 2, which both carry two *CSLD*s as tandem repeats. The tandem duplication may have occurred after WDG2 on the common ancestor of chromosomes 1 and 2 or before WGD1 followed by loss of one duplicate from the common ancestor of chromosomes 14 and 10/17. There is no evidence of loss following the WGD2. Intron structure is most parsimoniously explained by intron loss. CSLD 2 and 6 have three introns, the second of which is shared with *P. patens CESA*s (1). This intron 2 is present in *CSLD3* and *7*, but not *CSLD1*, *4*, *5* and *8* (indicated in red). Gain of intron 2 in CSLD2, 3, 6 and 7 is unlikely given that it is homologous with an intron in *P. patens CESA*s. It is possible that intron 2 was lost before WGD2 in the common ancestor of *CSLD5* and *8* and lost independently in the common ancestor of *CSLD1* and *4* before WGD2, but after tandem duplication of the common ancestor of *CSLD1, 3, 4* and *7*. Alternatively, tandem duplication and loss of intron 2 in the common ancestor of *CSLD1, 4, 5* and *8* may have occurred before WGD1 with loss of the paralog of the *CSLD3* and *7* common ancestor occurring before WGD2.

**Figure S3:**
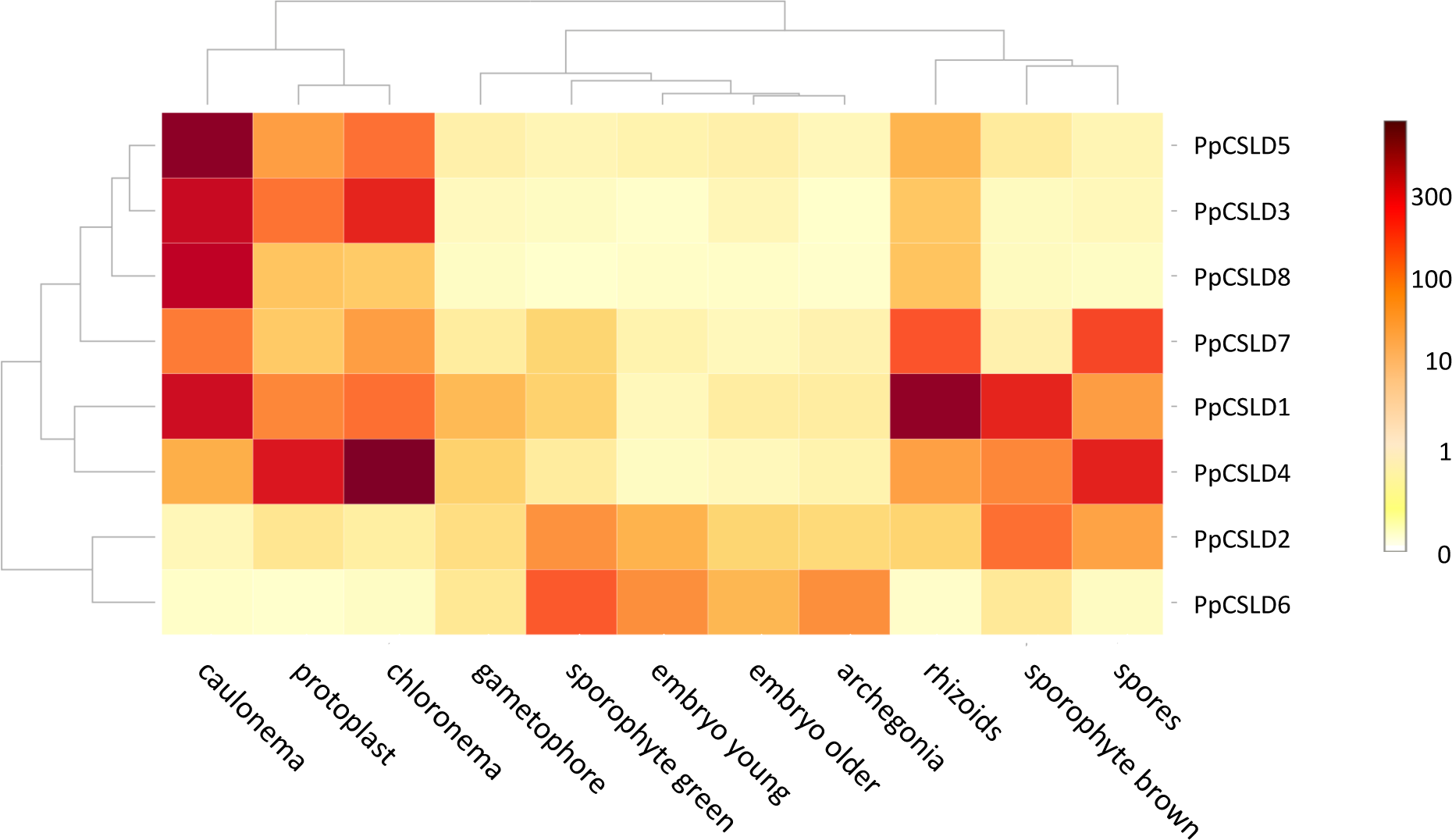
CSLD2 and CSLD6 transcripts are enriched in gametophores. Transcriptional profile of *P. patens CSLD*s at different developmental stages using a NimbleGene custom microarray (9) accessed from PEATmoss (10). *CSLD2* and CSLD6 had higher expression in gametophores and sporophytes compared to protonemal tissues (chloronema and caulonema). In contrast, the other six *P. patens CSLD*s were more highly expressed in protonemal tissues and had low expression in gametophores. Results from transcriptional profiling of *P. patens* developmental stages using a CombiMatrix array (11, 12) or RNA-seq (13) were generally consistent, although *CSLD2* transcripts were not detected in the RNA-seq analysis.

**Figure S4:**
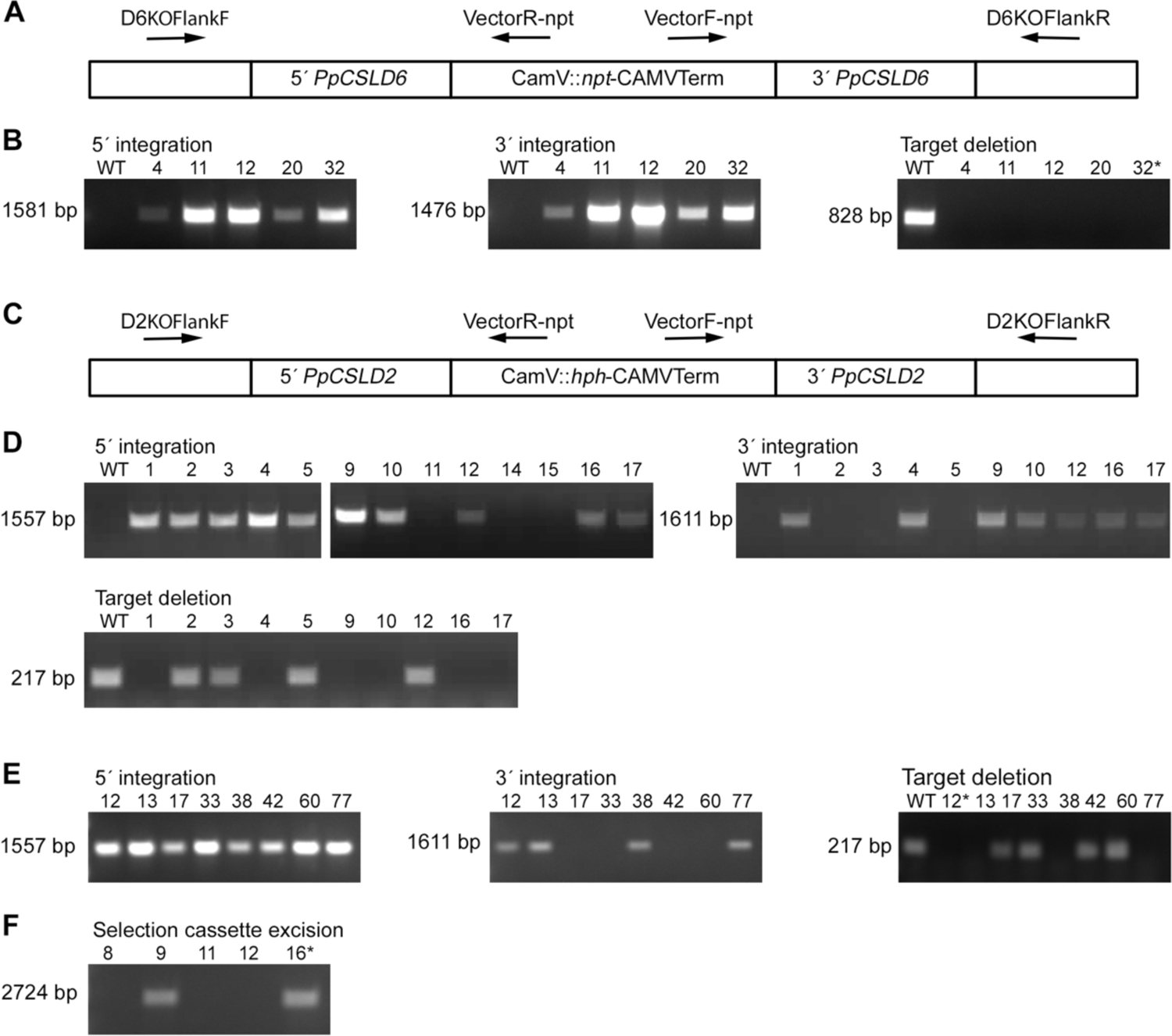
PCR-based genotyping of *ppcsld6*KO, *ppcsld2*KO and *ppcsld2/6*KO lines. (A) Primers used for amplification of the 5’ and 3’ integration sites are indicated as arrows on the diagram showing the PpCSLD6KO-npt vector integrated so as to delete *PpCSLD6*. (B) 5’ integration tested with primer pair D6KOFlankF/VectorR-npt produced the expected 1581 bp fragment in lines *csld6*KO-4, −11, −12, −20 and −32. 3’ integration tested with primer pair VectorF-npt/D6KOFlankR produced the expected 1476 bp fragment in the same 5 lines. Target deletion is verified in the *csld6*KO lines by the absence of a product from primers D6TargetF/D6TargetR, which anneal within the *CSLD6* coding sequence and amplify a 828 bp fragment in the wild type. (C) Primers used for amplification of the 5’ and 3’ integration sites are indicated as arrows on the diagram showing the PpCSLD2KO-hph vector integrated so as to delete *PpCSLD2*. (D) 5’ integration tested with primer pair D2KOFlankF/VectorR-hph produced the expected 1557 bp fragment in 10 *csld2*KO lines. 3’ integration tested with primer pair VectorF-hph/D2KOFlankR produced the expected 1611 bp fragment in 7 of those lines. Target deletion is verified in lines *csld2*KO-1, −4, −9, −10, −16 and −17 by the absence of a product from primers D2TargetF/D2TargetR, which anneal within the *CSLD2* coding sequence and amplify a 215 bp fragment in the wild type. (E) 5’ integration tested with primer pair D2KOFlankF/VectorR-hph produced the expected 1557 bp fragment in 8 *csld2/6*KO lines. 3’ integration tested by PCR with primer pair VectorF-hph/D2KOFlankR produced the expected 1611 bp fragment in 4 of those lines. Target deletion is verified in lines *csld2/6*KO-12, −13, −38 and −77 by the absence of a product from primers D2TargetF/D2TargetR, which anneal within the *CSLD2* coding sequence and amplify a 217 bp fragment in the wild type. (F) *cre*-mediated deletion of the selection cassette was verified for two csld2/6KO lines by amplification across the deletion site with primers D2KOFlankF/D2KOFlankR (2724 bp).

**Figure S5:**
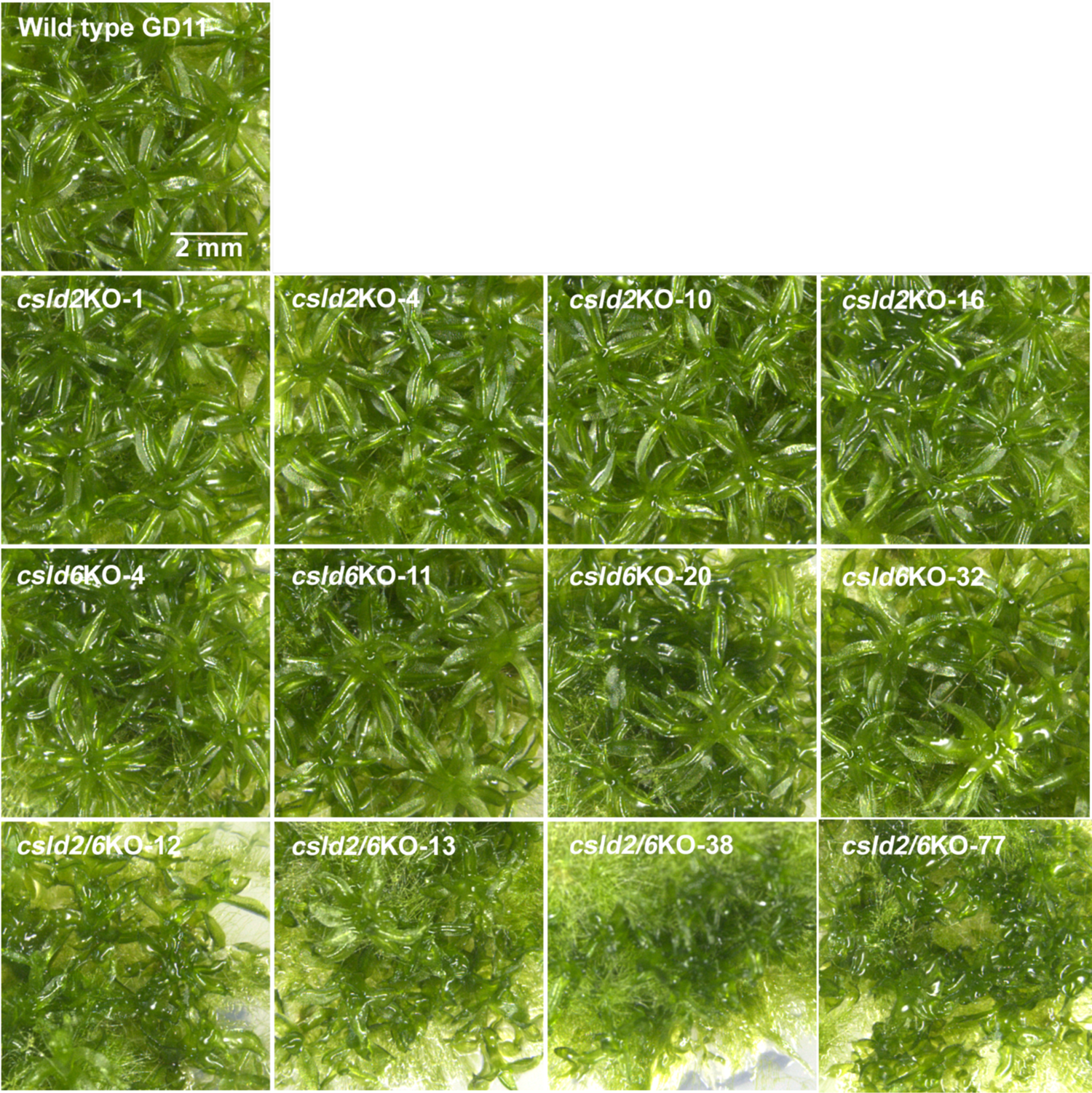
*CSLD2* and *CSLD6* are redundant and required for normal gametophore development. Gametophores from four independent lines each of *csld2*KO and *csld6*KO produce gametophores with wild type morphology. Four independent *csld2/6*KO lines produced gametophores with deformed phyllids.

**Figure S6:**
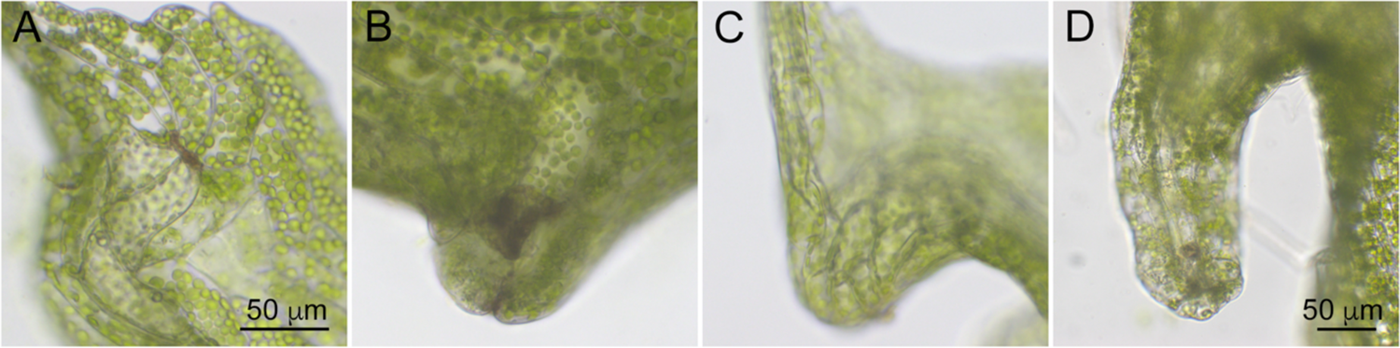
Tube structures form on *csld2/6*KO phyllids by alteration in the direction of cell expansion. (A) Phyllid cells surrounding cell separations alter their direction of cell elongation such that their long axes radiate from the separation. (B) Continued elongation of cells surrounding the separation results in formation of a bulge on the abaxial surface of the phyllid with the separation at the apex. (C) Cell division and expansion enlarges the bulge. (D) Continued elongation of the cells within the bulge forms a tube that protrudes from the abaxial surface of the phyllid.

**Figure S7:**
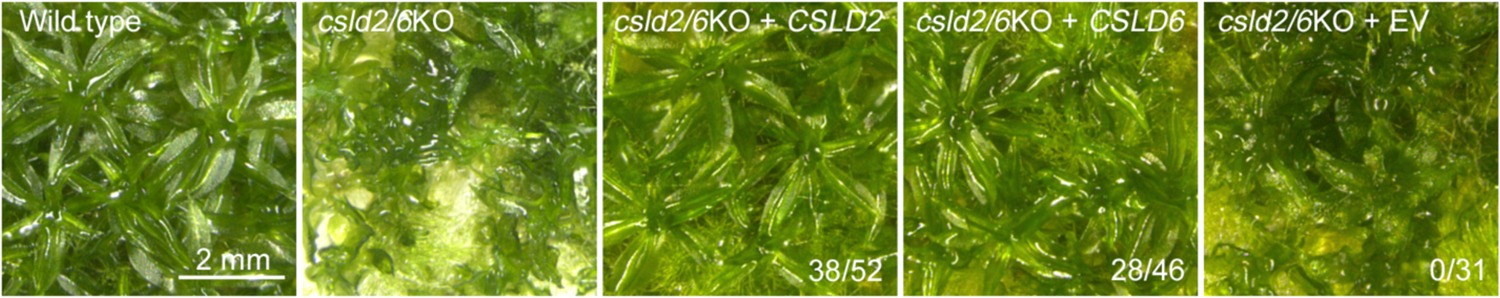
CSLD2 or CSLD6 rescues phyllid development defects. Wild type leaf morphology is restored when *csld2/6*KO plants are transformed with either a *CSLD2* or a *CSLD6* expression vector, but not an empty control vector (EV). Ratios indicate the number of transformed lines with normal gametophores over the total number of transformed lines with gametophores.

**Figure S8:**
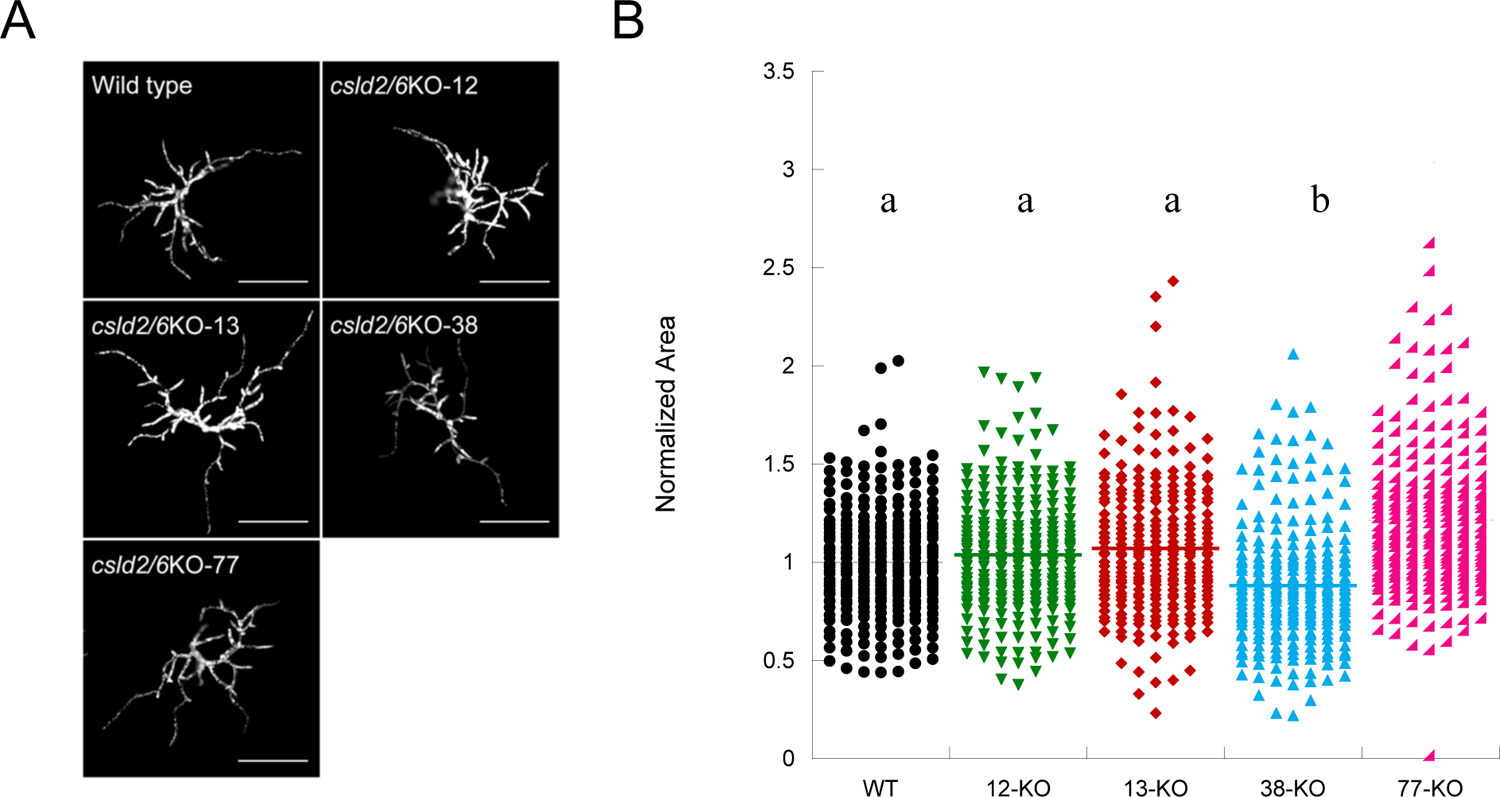
*CSLD2* and *CSLD6* are not required for protonemal development. (A) Chlorophyll autofluorescence images of 7-day old wild type and *csld2/6*KO plants regenerated from protoplasts. Scale bar, 250 µm. (B) Quantification of chlorophyll autofluorescence as a proxy for total plant area. For each of two experiments, 25 colonies were measured from each of 6 replicate plates for each genetic line. Area was normalized to the wildtype parent line. Significant differences determined by a one-way ANOVA analysis with a Tukey post hoc test (alpha = 0.05) are indicated by different letters.

**Figure S9:**
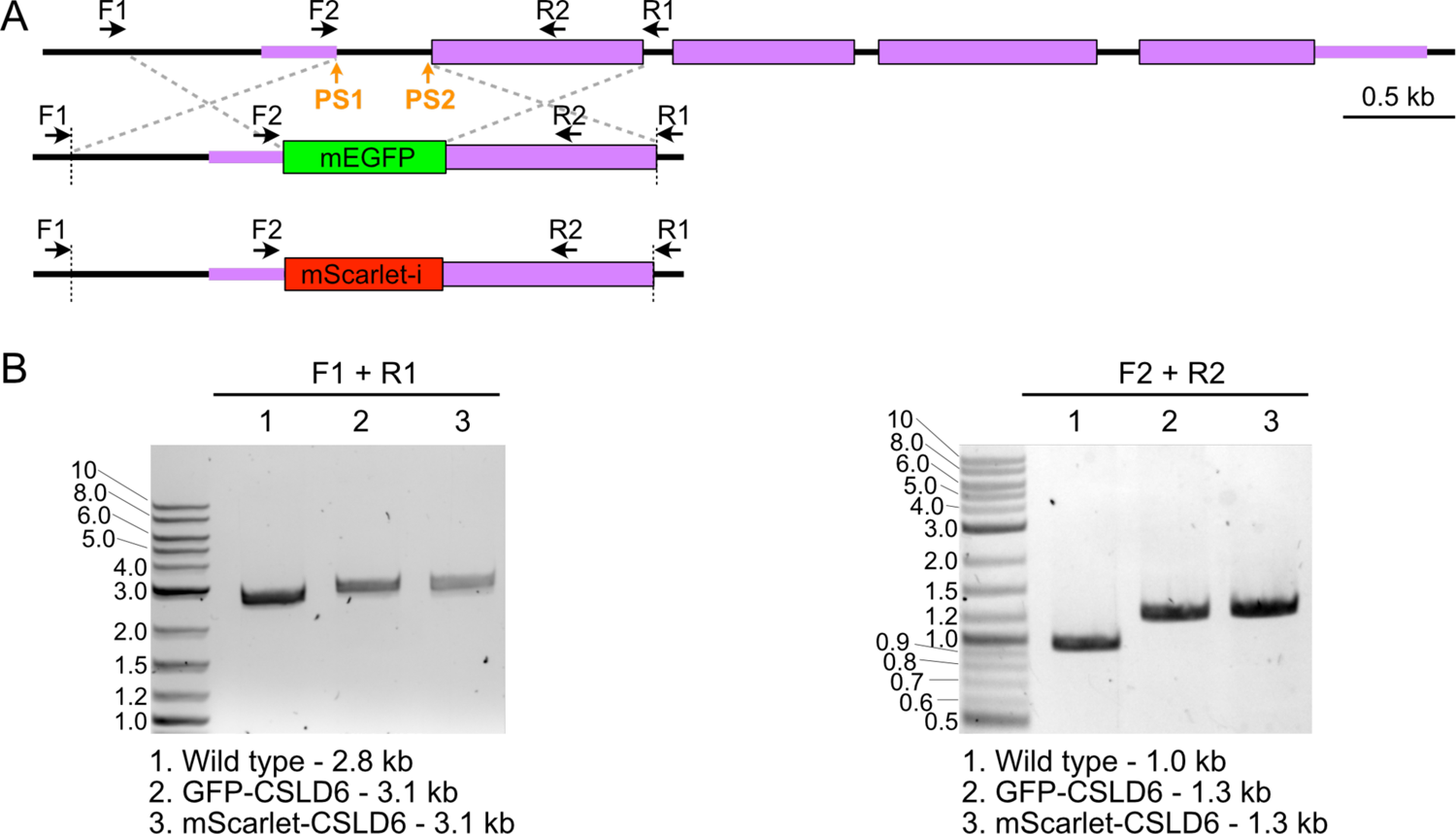
Molecular characterization of the tagged *CSLD6* locus. (A) Diagram illustrates the result of HDR mediated insertion of mEGFP (middle) or mScarlet-i (bottom) sequences in the *CSLD6* genomic locus. Coding exons are indicated by thick boxes and untranslated exons are indicated by thin boxes. Thin lines indicate intronic regions. Inserted sequences are denoted by thick colored boxes (green, mEGFP; red, mScarlet-i). The dashed vertical lines indicate the junction between the knock-in construct and upstream and downstream genomic sequences. Small arrows above the diagrams represent primers used for genotyping. Scale bar is 0.5 kb. (B) PCR products obtained with the indicated primer pairs using template DNA isolated from the indicated moss lines were separated on an agarose gel and stained with ethidium bromide. Molecular weight is indicated in kb. Predicted sizes for correct products are indicated below each gel.

**Figure S10:**
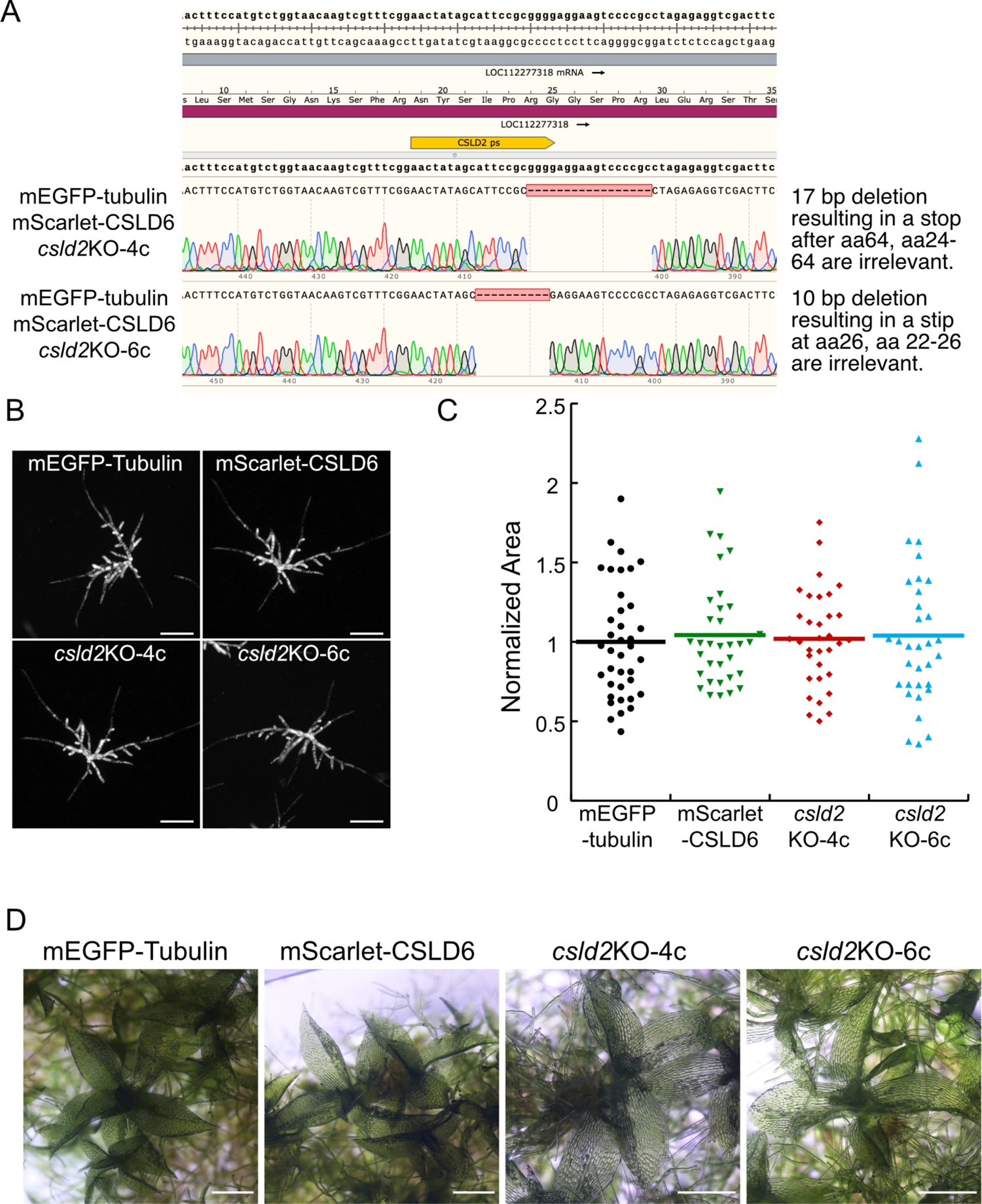
mScarlet-CSLD6 is functional. CRISPR-Cas9 mediated genome editing was used to generate *csld2*KO mutants in the mEGFP-tubulin/mScarlet-CSLD6 line. (A) Sequencing of two *csld2*KO alleles reveals CRISPR-Cas9 mediated genomic lesions. Zoom in of the region of the CSLD2 gene targeted by the protospacer (yellow bar). Genomic sequence is shown above the grey bar and predicted coding sequence is shown above the purple bar. Sanger sequencing results are shown below for each allele. (B) Chlorophyll autofluorescence images of 7-day old plants regenerated from protoplasts. Scale bar, 250 µm. (C) Quantification of chlorophyll autofluorescence as a proxy for total plant area. Area was normalized to the parent line, mEGFP-tubulin. There was no significant difference between the four groups as determined by a one-way ANOVA analysis with a Tukey post hoc test (alpha = 0.05). (D) Gametophore phyllids form normally in 3-4 week-old plants regenerated from ground tissue, demonstrating that the tagged CSLD6 is functional. Scale bar, 500 µm.

**Figure S11:**
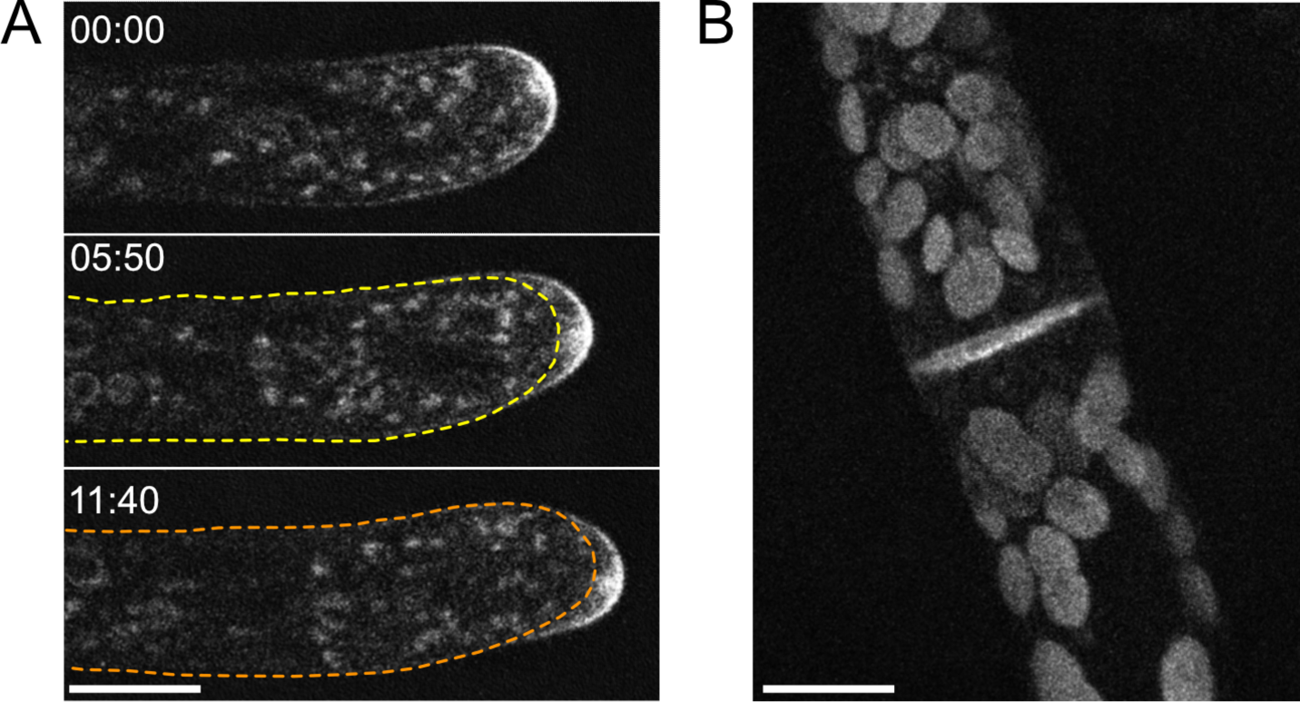
mEGFP-CSLD6 localization in protonemata. (A) Similar to mScarlet-CSLD6, mEGFP-CSLD6 is enriched in cytosolic punctae and at the apical plasma membrane of tip growing cells. Images are from a time lapse acquisition of confocal images of the medial plane of a growing protonemal cell. Time is indicated by min:sec. Yellow and orange dotted lines indicate the shape of the cell at 00:00, and 05:50 times, respectively. Scale bar, 10 µm. (B) Maximum projection of a confocal Z-stack of a dividing protonemal cell shows that mEGFP-CSLD6 accumulates at the cell division plane. Large globular structures are chloroplasts which are more concentrated near the division plane and auto-fluoresce in the GFP channel.

**Figure S12:**
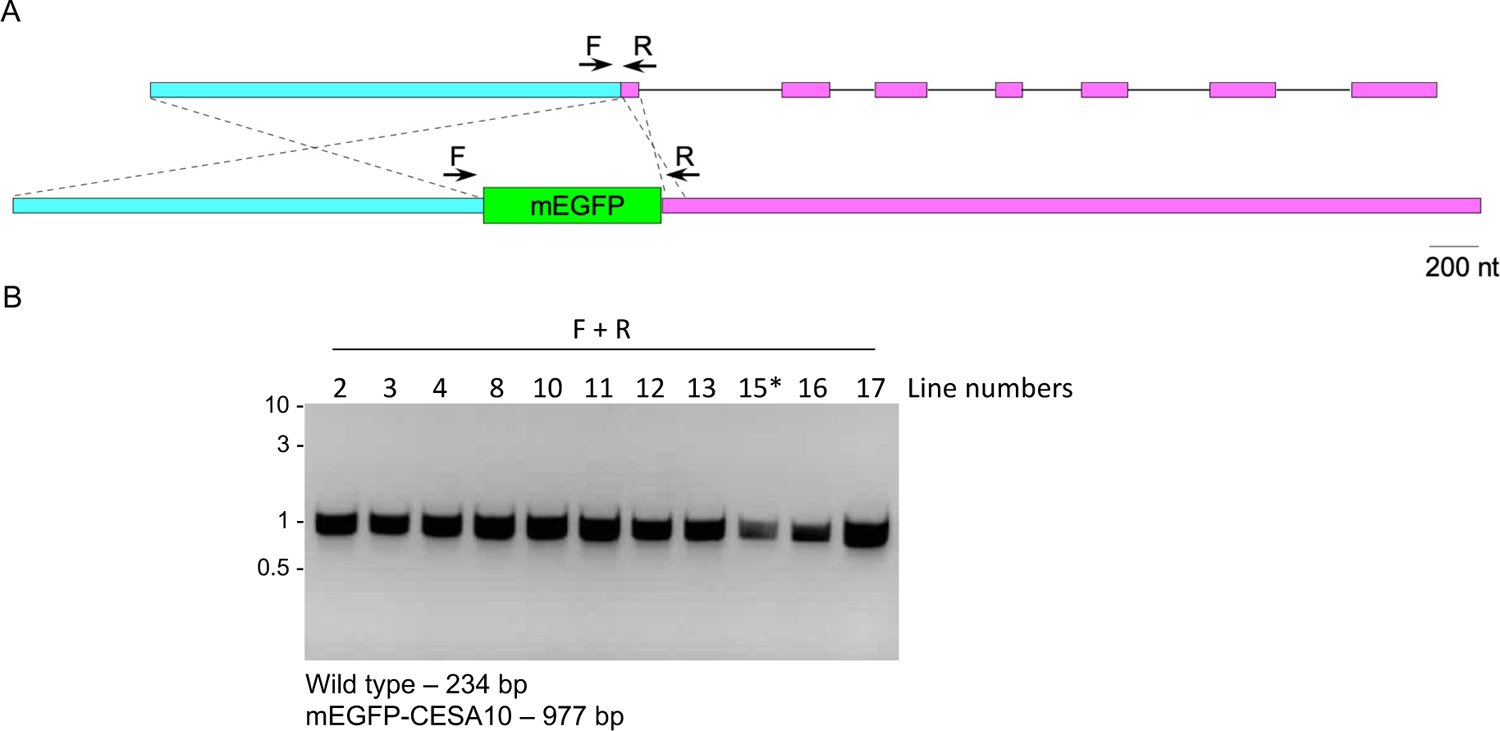
Molecular characterization of the tagged *CESA10* locus. (A) Diagram illustrates the result of HDR mediated insertion of mEGFP sequences from the homology repair plasmid (bottom) into the *CESA10* genomic locus (top). Exons (first 7 shown) are indicated by pink boxes and the cloned promoter is indicated by cyan boxes. Thin lines indicate intronic regions. The inserted mEGFP sequence is denoted by a thick green box. Small arrows above the diagrams represent primers used for genotyping. (B) PCR products obtained with primer pairs using template DNA isolated from the indicated moss lines were separated on an agarose gel and stained with ethidium bromide. The asterisk indicates the line chosen for imaging after sequencing the PCR product. Molecular weight is indicated in kb. Predicted sizes for correct products are indicated below the gel.

**Figure S13:**
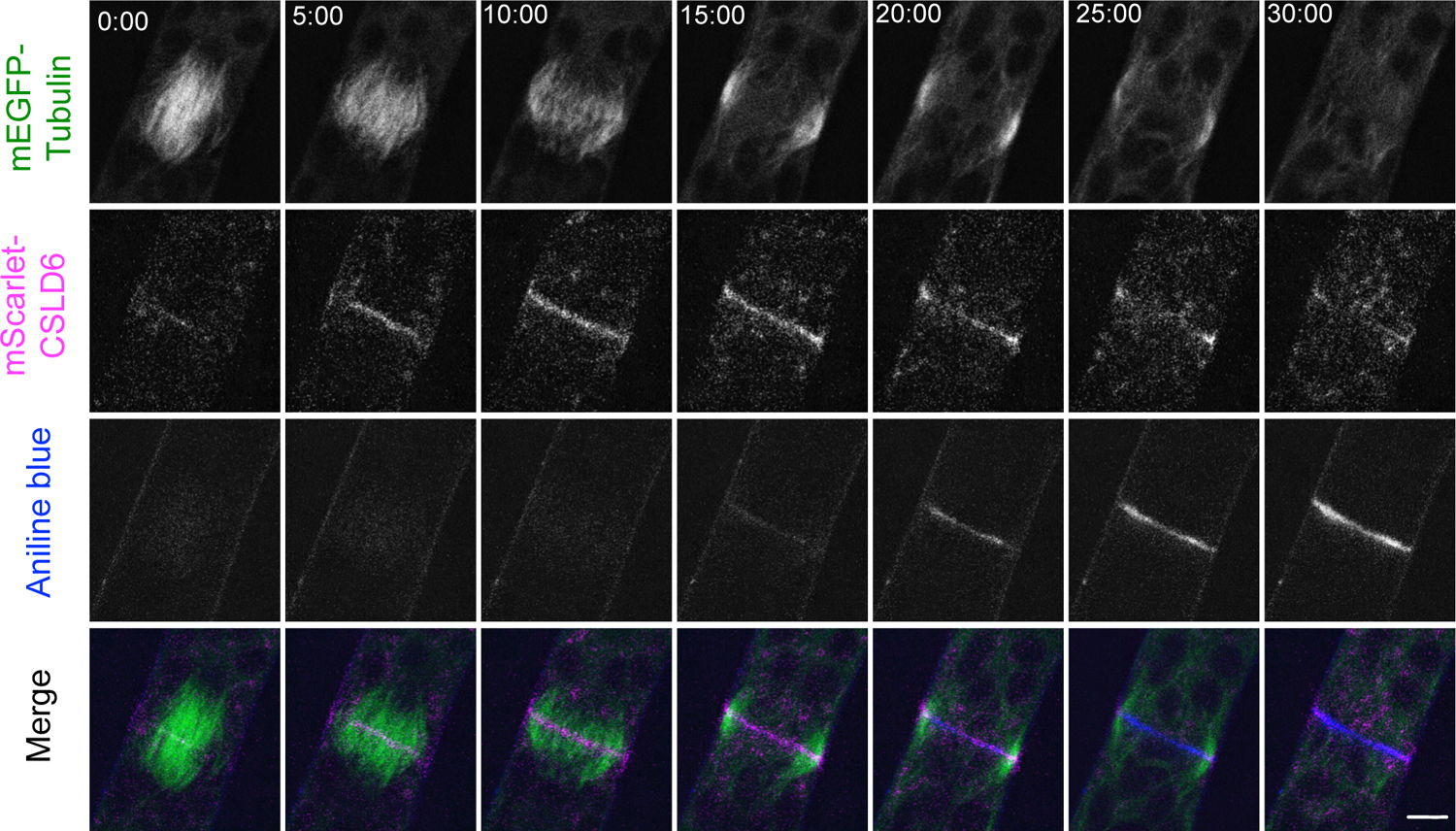
Isoxaben does not affect cell plate formation. Cell division in moss protonemata expressing mScarlet-CSLD6 (magenta in merge) and mEGFP-tubulin (green in merge), stained with aniline blue for callose (blue in merge). Cell treated with 20 µM isoxaben. Images are single focal plane confocal images from a time-lapse acquisition. Scale bar, 5 µm. Time stamp, min:sec.

**Figure S14:**
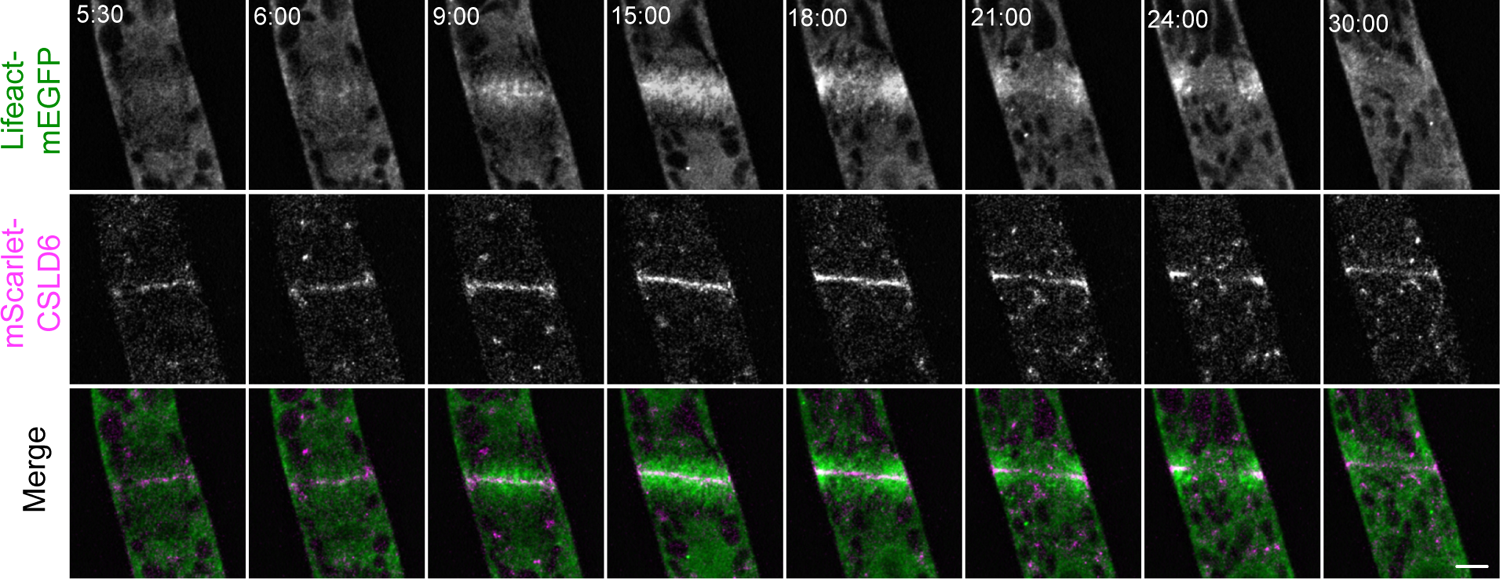
CSLD6 accumulates in the nascent cell plate slightly earlier than actin. Moss protonemata expressing lifeact-mEGFP (green in merge) and mScarlet-CSLD6 (magenta in merge). CSLD6 appears in the cell plate (5:30) earlier than actin (6:00). Images are single focal plane confocal images from a time-lapse acquisition. Scale bar, 5 µm. Time stamps, min:sec.

**Figure S15:**
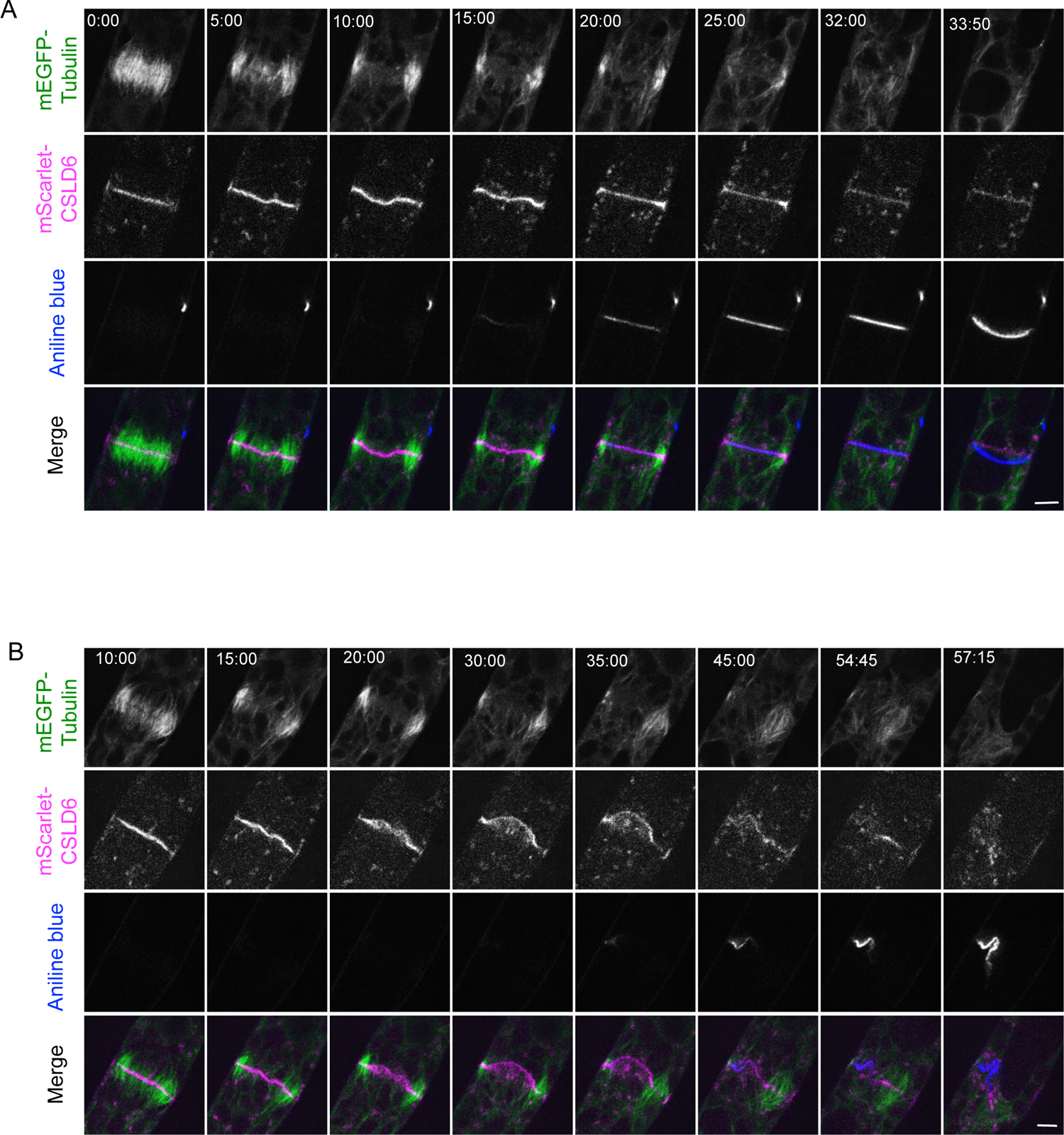
Additional examples demonstrating that CSLD activity and actin stabilize the nascent cell plate. Cell divisions in moss protonemata expressing mScarlet-CSLD6 (magenta in merge) and mEGFP-tubulin (green in merge), stained with aniline blue for callose (blue in merge). (A) Cell treated with 10µM DCB. The cell plate buckles (5:00-15:00) but straightens again afterwards (20:00-32:00). (B) Cell treated with 25 µM LatB and 10 µM DCB. The cell plate buckles and remains deformed even after callose deposition. Images are single focal plane confocal images from time-lapse acquisitions. Scale bars, 5 µm. Time stamps, min:sec.

